# No leading-edge effect in North Atlantic harbor porpoises: Evolutionary and conservation implications

**DOI:** 10.1101/2020.11.03.366542

**Authors:** Yacine Ben Chehida, Roisin Loughnane, Julie Thumloup, Kristin Kaschner, Cristina Garilao, Patricia E. Rosel, Michael C. Fontaine

## Abstract

Understanding a species response to past environmental changes can help forecast how they will cope with ongoing climate changes. Harbor porpoises are widely distributed in the North Atlantic and were deeply impacted by the Pleistocene changes with the split of three sub-species. Despite major impacts of fisheries on natural populations, little is known about population connectivity and dispersal, how they reacted to the Pleistocene changes and how they will evolve in the future. Here, we used phylogenetics, population genetics, and predictive habitat modelling to investigate population structure and phylogeographic history of the North Atlantic porpoises. A total of 925 porpoises were characterized at 10 microsatellite loci and one-quarter of the mitogenome (mtDNA). A highly divergent mtDNA lineage was uncovered in one porpoise off Western Greenland, suggesting that a cryptic group may occur and could belong to a recently discovered mesopelagic ecotype off Greenland. Aside from it and the southern sub-species, spatial genetic variation showed that porpoises from both sides of the North Atlantic form a continuous system belonging to the same subspecies (*Phocoena phocoena phoceona*). Yet, we identified important departures from random mating and restricted intergenerational dispersal forming a highly significant isolation-by-distance (IBD) at both mtDNA and nuclear markers. A ten times stronger IBD at mtDNA compared to nuclear loci supported previous evidence of female philopatry. Together with the lack of spatial trends in genetic diversity, this IBD suggests that migration-drift equilibrium has been reached, erasing any genetic signal of a leading-edge effect that accompanied the predicted recolonization of the northern habitats freed from Pleistocene ice. These results illuminate the processes shaping porpoise population structure and provide a framework for designing conservation strategies and forecasting future population evolution.

## 1 Introduction

Movements of highly mobile marine species are potentially unrestricted over vast geographical distances. Absence of evident barriers to gene flow challenges, at least in theory, the process of population divergence (Palumbi, 1994). High dispersal ability should favor population homogeneity and limit spatial genetic structure over large geographic scales (Palumbi, 1994). Yet, highly dispersive species like cetaceans exhibit strong population structure, even at small geographical scales (Hoelzel, 2009; Vachon et al., 2018). These patterns are usually driven by both current and historical mechanisms. Oceanographic features, ecological specialization, and complex behaviors are often invoked as mechanisms contributing to limit dispersal and structure cetacean populations (Fontaine et al., 2007; Foote et al., 2016; Hoelzel, 2009; Vachon et al., 2018). Historical environmental variation during the Quaternary glaciations had a major role in shaping patterns of genetic structure and diversity in both terrestrial and marine environments (Hewitt, 2000). Habitat releases during post-glacial periods have opened ecological niches and created ecological opportunities that spurred the evolution of cetaceans (Slater et al., 2010; Steeman et al., 2009). In particular, major glaciations of the Last Glacial Maximum (LGM) compressed suitable habitats towards the equator. The successional cold and warm periods resulted in large habitat contraction and expansion. These environmental fluctuations promoted population subdivision and the formation of sub-species, as observed in multiple cetacean species like killer whales *Orcinus orca* (Morin et al., 2015), bottlenose dolphins *Tursiops truncatus* (Louis et al., 2014; Louis et al., 2020; Moura et al., 2020), and harbor porpoises *Phocoena phocoena* (Fontaine, 2016; Fontaine et al., 2014; 2010; Rosel et al., 1995; Tolley & Rosel, 2006). These historical demographic events leave a detectable imprint on genetic variation, which can be used to identify divergent lineages, reconstruct species evolutionary history, unravel the impacts of past climatic processes on the current spatial distributions of species, and formulate hypotheses about population and species future evolution (Hewitt, 2000; Hickerson et al., 2010).

Harbor porpoises are amongst the smallest cetacean species. This species has been described as ‘*living in the fast lane*’ due to its life history traits marked by high reproductive demands, short generation time (∼10 years per generation) and relatively short life span for a cetacean (∼12 years and up to 24 years) (Lockyer, 2007; Read & Hohn, 1995). Porpoises must thus rely on a regular food supply to meet their metabolic demands (Hoekendijk et al., 2018; Wisniewska et al., 2016). They are opportunistic feeders, feeding mostly on the continental shelf, often targeting demersal or benthic species (ex. Santos & Pierce, 2003; but see Nielsen et al. 2018). Coined the ‘aquatics shrews’ of the sea, prey availability has been shown to be an important driver of porpoise movements (Johnston et al., 2005; Sveegaard et al., 2012; Wisniewska et al., 2016) and local densities (Hammond et al., 2013; Marubini et al., 2009; Waggitt et al., 2018). These animals are thus expected to be highly susceptible to environmental changes, increasing sea temperature, and prey displacements or modifications (Lambert et al., 2014; MacLeod et al., 2005). Furthermore, populations are heavily impacted by incidental catches in commercial fisheries (Braulik et al., 2020; ICES WGBYC 2019; NAMMCO & IMR, 2019; Stenson, 2003). In order to predict future movements and distribution in a changing environment and the impact of heavy bycatch casualties on natural populations, we must understand how populations are genetically structured and connected to each other, and how they have reacted to past environmental changes.

Distributed in subpolar to temperate waters of the Northern Hemisphere (Fontaine, 2016; Gaskin, 1984; Read, 1999), harbor porpoises are mostly found in coastal waters of the North Pacific, North Atlantic, and in the Black Sea, with three subspecies currently officially recognized: *P. p. vomerina, P. p. phocoena*, and *P. p. relicta*, respectively. Porpoises from the upwelling zones off Iberia and Mauritania have been recently identified as genetically divergent as *P. p. phocoena* and *P. p. relicta*, based on DNA sequence analysis of a quarter of the mitogenome (Fontaine, 2016; Fontaine et al., 2007; 2014). They have thus been proposed to belong to a separate subspecies (formally unnamed subspecies and possibly *P. p. meridionalis* as suggested in Fontaine et al. 2014) due to their distinctiveness in terms of genetics, morphology and ecology (Fontaine, 2016; Fontaine et al., 2007; 2014). As a formal description has not yet been made for this subspecies, we refer in this paper to these porpoises as the Iberia-Mauritania porpoises (*IBMA*). Demo-genetic inferences suggested that the three lineages in the North Atlantic and the Black Sea split during the LGM and were following independent evolutionary trajectories making them distinct evolutionary significant units (Fontaine, 2016; Fontaine et al., 2014; 2010). These lineages originated from an initial split of ancestral populations stemming from the North Atlantic colonizing the Mediterranean Sea. *P. p. relicta* and *IBMA* likely descended from these ancestral populations that inhabited the Mediterranean Sea during cold and nutrient-rich periods prevailing during the LGM. More recently, *IBMA* and *P. p. phocoena* populations in the North Atlantic likely came back into contact establishing a contact zone in the northern part of the Bay of Biscay during postglacial warming (Fontaine et al., 2014; 2017). This hybridization zone is characterized by strong habitat differences in terms of oceanographic conditions compared to the prevailing cold and highly productive waters found to the north on the European continental shelf or south along the Iberian coast.

While the evolution of the porpoises surrounding the Mediterranean Sea has been fairly well studied (Alfonsi et al., 2012; Fontaine et al., 2007; 2014; 2012; 2010; Tolley & Rosel, 2006; Viaud-Martinez et al., 2007; see the review of Fontaine, 2016), the phylogeographic history of *P. p. phocoena* spreading north of the Bay of Biscay on both sides of the North Atlantic remains under debate. This subspecies is fairly continuously distributed from the French Biscayan waters northward to the North and Barents Seas, and westward across the North Atlantic, around the Faroe Islands, Iceland and West Greenland and then south along Western North Atlantic shorelines of Canada and eastern coast of the

United States (Fontaine, 2016; Gaskin, 1984; Read, 1999). It has been hypothesized (*hypothesis 1*) that porpoises on each side of the North Atlantic could have evolved independently, recolonizing their current ranges from distinct source populations living in distinct southern refugia during the LGM (Gaskin, 1984; Rosel et al., 1999b; Yurick & Gaskin, 1987). Management plans by the IWC in 1996 (Donovan & Bjørge, 1995) considered them as separate evolutionary entities based on this suggestion. However, another hypothesis (*hypothesis 2*) could be that suitable habitats were shifted southward without any loss of connectivity between populations from each side of the North Atlantic, therefore leading to no lineage split between each side of the North Atlantic and to limited population contraction. These two hypotheses are expected to leave distinct signatures on the genetic variation of natural populations. Under *hypothesis 1* (two distinct refugia), two divergent genetic lineages would be expected, one on each side of the North Atlantic, similar to what was previously observed around the Mediterranean Sea (Fontaine, 2016; Fontaine et al., 2014). Furthermore, a gradient in genetic diversity would be expected, with higher diversity in southern areas where populations would have survived since the LGM and genetic diversity decreasing northward toward the most recently colonized northern temperate habitats. This would be consistent with a leading-edge colonization effect where allele surfing leads to gradual loss of diversity due to genetic drift associated with serial founder effects toward the colonization front (Excoffier et al., 2009; Excoffier & Ray, 2008). Under *hypothesis 2*, we would not expect any distinct lineages in either side of the North Atlantic, but rather a single lineage with relatively high diversity. Genetic diversity would be close to a migration-drift equilibrium, characterized by more homogeneous spatial patterns of genetic diversity, possibly with evidence of isolation-by-distance if intergenerational dispersal is spatially restricted (Hutchison & Templeton, 1999). Variation in local population density could also be expected, decreasing toward southern habitats where warmer waters would become less suitable for a cold-water adapted species like harbor porpoises. No study combining phylogeographic approaches together with habitat modelling have been conducted to date to tease apart these hypotheses.

Previous genetic studies based on short fragments (∼400 base-pairs, bps) of the mitochondrial control region harbored limited phylogenetic information, as shown by the previous shallow and poorly resolved phylogenetic trees and networks (Rosel et al., 1999b; Tolley & Rosel, 2006; Viaud-Martinez et al., 2007; Wang et al., 1996). For example, with such a short fragment, *IBMA* porpoises from Iberian and Mauritanian waters could be identified as genetically differentiated from the other subspecies in terms of haplotype frequencies, but the full extent of their divergence only became clear when analyzing fragments ten times longer covering one quarter of the mitogenome (Fontaine et al., 2014). In the present study, we revisited the population genetic structure and phylogeography of harbor porpoises across the entire North Atlantic distribution range. Combining phylogenetic and spatial population genetic approaches together with predictive habitat modelling, we tested the two hypotheses of post-glacial evolution described above. For this purpose, we reanalyzed samples from the North West Atlantic (NWA) waters previously used in Rosel et al. (1999a) with similar genetic markers as those used in Fontaine et al. (2014). These included sequences from one-quarter of the mitogenome and ten highly polymorphic microsatellite loci. We combined these new data from NWA porpoises with those from the central and Eastern North Atlantic (NEA) from Fontaine et al. (2014), which included samples collected during the same time period.

Specifically, we (1) assessed whether distinct mtDNA lineages were present in *P. p. phocena*, possibly indicating distinct glacial refugia in the North Atlantic during the LGM; (2) evaluated the postglacial population responses and recolonization routes from the analyses of spatial patterns of genetic variation and whether porpoises recolonized their present range from one or multiple refugia; (3) assessed the impact of environmental changes on the distribution of harbor porpoise by modelling the evolution of suitable habitats at present, during the LGM, and by the year 2050; and (4) analyzed dispersal behaviors at the North Atlantic scale and whether restricted dispersal could generate isolation-by-distance patterns as previously reported in NEA (Fontaine et al., 2007) and NWA (Rosel et al., 1999a; Wang et al., 1996). Moreover, we investigated the extent of sex-biased dispersal with strong female philopatry previously suggested in the species (Rosel et al., 1999a; Wang et al., 1996). Here, we reassessed this effect at the scale of the entire North Atlantic distribution of the *P. p. phocoena* subspecies by comparing genetic markers with contrasted inheritance modes (bi-parentally inherited microsatellite vs maternally inherited mtDNA).

## 2 Material and methods

### 2.1 Sampling and data collection

We combined previous nuclear microsatellite genotype data at ten loci for the 768 samples from Central and Eastern North Atlantic populations (Fontaine et al., 2014) with 173 newly genotyped samples from NWA. We used the samples from Rosel et al. (1999a) and genotyped them with the same markers as in Fontaine et al. (2014). The NWA sampling was from a same time cohort (1990-1999) as those analyzed in Fontaine et al. (2014) and included 29 individuals from West Greenland (WGLD), 60 from Canada (CA) and 84 from the United States (US) collected from incidentally entangled and stranded animals (Table S1).

Total genomic DNA was extracted from the skin tissues using PureGene and DNeasy Tissue kits (Qiagen) following the manufacturer’s recommendations. DNA quality and quantity of the extraction products were checked by electrophoresis on an agarose gel stained with ethidium bromide and using a fluorometric quantitation procedure using a Qubit v.3.0 (Thermo Scientific). The microsatellite genotyping procedure followed the protocol described in Fontaine et al. (2007; 2006). All genotypes were checked visually and individuals with missing data or ambiguous alleles were genotyped twice or three times. Only individuals with at least 6 successfully genotyped loci were kept for downstream analyses. For various analyses, we subdivided the samples into ten putative subpopulations (hereafter called geographical regions) (Figure 1, Table S1): Black Sea (BS), Mauritania (MA), Iberia (IB), northern Bay of Biscay (NBB), North Sea (NS), North Norway (NN), Iceland (IC), Greenland (WGLD), Canada (CA) and the United States (US). For analyses requiring fine-grained geographic partitioning for microsatellite data, we subdivided further geographical regions into 30 local geographical sub-groups (Figure S1 and Table S2).

**Figure 1.**
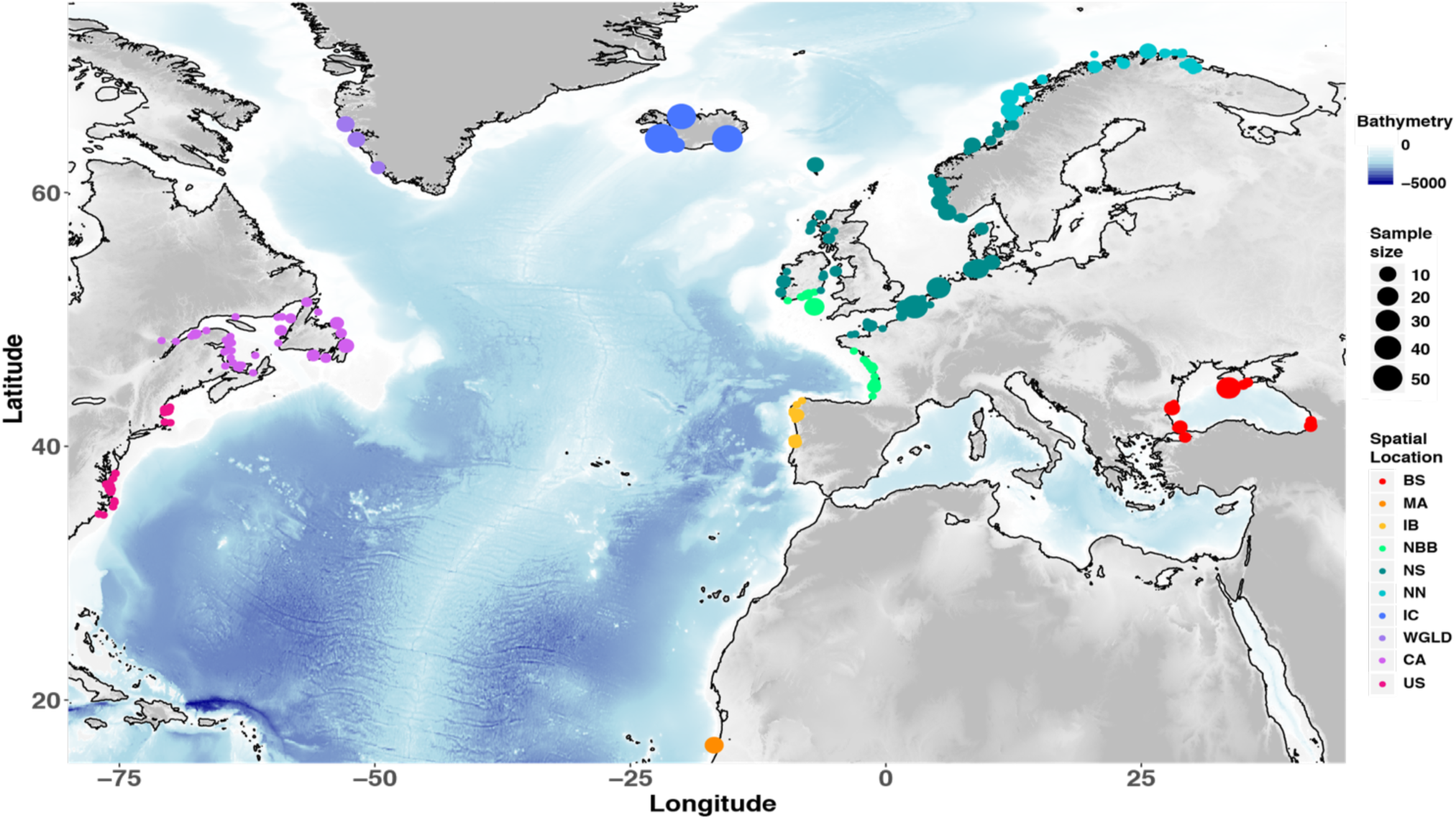
Map showing the sampling locations of individual porpoises in the North Atlantic and the 10 genetic groups defined as geographical regions in this study (see Figure S1 for finer delimitations into 30 subgroups). Sampling locations are based on approximate GPS coordinates or reported discovery location. Acronyms are as follow: BS=Black Sea; MA=Mauritania; IB=Iberia: NBB=North Bay of Biscay; NS=North Sea; NN=North Norway; IC=Iceland; WGLD=West Greenland; CA=Canada; US=United States.

In addition to autosomal microsatellite data, a 4,465 bp fragment of the mtDNA genome encompassing five coding regions (*CytB, ATP6, ATP8, ND5*, and *COXI*) was obtained for a subsampling of 55 individuals from NWA (Table S1), including also five new samples from Iceland, following the PCR amplification protocol described previously (Fontaine et al., 2014). PCR products were visualized under UV light before being prepared for Sanger sequencing on a 1% agarose gel stained with ethidium bromide. Purification of the PCR products used ExoSAP-IT™ PCR Product Cleanup Reagent (ThermoFisher Scientific) and products were then sent to GATC Biotech for sequencing. We visually inspected the quality of the sequence electropherograms and manually edited them using *Geneious* v.8.1.9 (Kearse et al., 2012). For each gene fragment, forward and reverse sequences were assembled into contigs and resulting contigs of each individual were visually inspected before making a multiple sequence alignment. We concatenated the five genes in accordance with previous data (Fontaine et al., 2014) following the same order (*ATP6-8, COI, Cyt-B, ND5*). The 55 newly sequenced NWA samples were combined with 81 previously generated mtDNA sequences from central and eastern North Atlantic porpoises (Fontaine et al., 2014). We added also 14 sequences from the North Pacific harbor porpoise *P. p. vomerina* subspecies from Ben Chehida et al. (2020) to the 136 North Atlantic sequences in order to place North Atlantic populations into a broader phylogeographic context (Table S3). Two sequences from the closest outgroup species, the Dall’s porpoise *Phocoenoides dalli*, were added from Fontaine et al. (2014) for the phylogenetic analyses. We performed the multiple sequence alignment using *MUSCLE* (Edgar, 2004) with default settings. MtDNA data were divided into the same 10 geographical regions as used for the microsatellite data for the data analyses (Figure 1, Table S1).

### 2.2 Mitochondrial phylogenetic relationships

We evaluated genetic relationships between mtDNA haplotypes using three different methods. First, we estimated phylogenetic relationships among unique haplotypes using the maximum-likelihood (ML) approach of *PHYML* (Guindon et al., 2010) implemented in *Geneious* v.8.1.9 (Kearse et al., 2012). Prior to phylogenetic reconstruction, we used *jModelTest* v.2.1 (Darriba et al., 2012) to identify the model of nucleotide evolution best fitting our dataset. The tree was rooted with two sequences from Dall’s porpoise. Node support was estimated using 1,000 bootstrap replicates. The resulting tree was visualized using *ggtree* v.1.4 (Yu et al., 2018) in the R statistical environment v.3.5.1 (R Core Team, 2018). In addition to the phylogenetic trees, we reconstructed a median-joining haplotype network using *PopART* (http://popart.otago.ac.nz), which allows displaying distances among haplotypes in a number of mutational steps and reticulations among them. Finally, we also summarized genetic relationships among haplotypes using a non-metric multidimensional scaling (nMDS) based on the Jukes-Cantor genetic distances between pairs of sequences. We used *MEGA* v.7.0 (Kumar et al., 2016) to calculate the genetic distance and the R package *ecodist* (Goslee & Urban, 2007) to compute the nMDS.

### 2.3 Population genetic structure and differentiation

We investigated the genetic structure among porpoises from Northwest (NWA: US, CA, WGLD) and Northeast (NEA: IC, NN, NS) Atlantic at the microsatellite and mitochondrial loci. For the microsatellite dataset, we first explored patterns of genetic variation and structure using multivariate methods, including a principal components analysis (PCA) (Patterson et al., 2006), a discriminant analysis of principal components (DAPC) (Jombart et al., 2010), and a spatial PCA (sPCA) (Jombart et al., 2008). The three approaches were performed using the *Adegenet* v2.1.1 R package (Jombart, 2008) on centered genetic data (i.e., set to a mean allele frequency of zero). The PCA was run with missing data replaced by the mean. The DAPC was performed without any missing data using the subspecies or geographical sub-groups as *a priori* groupings with the number of principal components set to 30 and 34, respectively, following alpha-score indication as recommended by the author. Additionally, we ran a DAPC only on *P. p. phocoena* individuals (NAT) maximizing the difference between NWA and NEA or among geographical subgroups in order to assess fine-scale population structure. Then, we displayed genetic variance with a spatial structure using a sPCA, which accounts for spatial autocorrelation among allele frequencies. We used the Delaunay triangulation as a connection network on a subset of 729 individuals with no missing data in both geographic coordinates and microsatellite genotypes. We used both ‘global’ and ‘local’ test procedures based on Monte Carlo permutations (10^4^ permutations) to interpret the significance of the spatial principal components in the sPCA (Jombart et al., 2008). ‘Global’ structure relates to patterns of spatial genetic structure, such as patches, clines, IBD and intermediates, whereas ‘local structure’ refers to strong differences between local neighborhoods.

We also used the model-based Bayesian clustering algorithm of *STRUCTURE* v.2.3.4 (Hubisz et al., 2009) to estimate individual genetic ancestry. *STRUCTURE* works by leveraging the fact that population structure induces departures from Hardy-Weinberg and linkage equilibrium (HWLE) expectations among loci. Therefore, contrary to multivariate methods, *STRUCTURE* identifies genetic clusters by minimizing departures from HWLE. It estimates the genetic ancestry proportions of each individual multilocus genotype to *K* ancestral clusters. *STRUCTURE* was performed for a data set with and without missing data. We used the *locprior* admixture model with correlated allele frequencies which is capable of detecting weak genetic structure when it exists without forcing it (Hubisz et al., 2009). The 30 geographical sub-group delimitations (Figure S1 and Table S2) were used to inform the *locprior* model. We tested different numbers of plausible clusters (*K*) ranging from 1 to 5. Each run used 1×10^6^ iterations after a burn-in of 1×10^5^ iterations. To evaluate the convergence of the Monte Carlo Markov Chains (MCMC), we performed 10 independent replicates for each *K* value and checked the consistency of the results using *CLUMPAK* v.1.0 (Kopelman et al., 2015). We determined the best *K* value using the log-likelihood of the data for each *K* value using *STRUCTURE HARVESTER* v.0.6 (Earl & vonHoldt, 2011) and by visually inspecting newly created clusters with increasing *K* values (Vercken et al., 2010). We plotted the results as barplots using *CLUMPAK*. Finally, we investigated geographical variation in genetic ancestry coefficients (Q) of the predominant solution for the best *K* by computing the spatial interpolation of Q values among sampling locations using the script provided at http://membres-timc.imag.fr/Olivier.Francois/TESS_Plot.html. We also ran *STRUCTURE* in a supervised way focusing only on *P. p. phocoena* individuals, as it has been suggested that rerunning *STRUCTURE* on identified clusters may reveal finer scale genetic structuring (Evanno et al., 2005).

Pairwise genetic differentiation among geographical regions and sub-groups at microsatellite loci was assessed using the *F*_*ST*_ estimator of Weir and Cockerham (1984) in the R package *diversity* v.1.9 (Keenan et al., 2013). The *F*_*ST*_ significance was tested using an exact test implemented in *GENEPOP* v.1.1.3 (Rousset, 2008) in R using 10,000 iterations. Geographical sub-groups with less than 10 individuals were excluded. We estimated the degree of mtDNA differentiation in terms of haplotype frequencies between pairwise geographical regions using the *F*_*ST*_ estimator of Hudson et al. (1992). Its significance level was tested using 1,000 permutations on the *Snn* statistics (Hudson, 2000). These calculations were conducted in *DnaSP* v.4.5 (Librado & Rozas, 2009). We also calculated the *Φ*_*ST*_ estimator (Excoffier et al., 1992) using *Arlequin* v3.5 (Excoffier & Lischer, 2010). *Φ*_*ST*_ exploits the degree of pairwise differentiation in haplotype frequencies together with nucleotide sequence divergence between pairs of geographical regions. *Φ*_*ST*_ values were computed assuming a TN93 substitution model (Tamura & Nei, 1993) and significance was assessed using 1,000 permutations. In addition to the two estimators of population differentiation, we also calculated the Jukes-Cantor based *d*_*A*_ distances (Nei, 1987) using *DnaSP* which provides a measure of net mtDNA divergence between pairs of groups.

### 2.4 Patterns of genetic diversity

We first tested linkage disequilibrium (LD) among microsatellite loci within putative groups using a *G*-test (Weir, 1996) implemented in *GENEPOP* in R (Rousset, 2008). We also quantified multi-locus LD within each geographical region using the *R*_*D*_ statistics (Agapow & Burt, 2001) and assessed its significance level using 1,000 permutations in *Poppr* v.2.8.3 (Kamvar et al., 2014). Within group departure from Hardy–Weinberg equilibrium was assessed using the exact tests implemented in *diveRsity* in R using 10,000 iterations and quantified it using the *F*_*IS*_ (Weir & Cockerham, 1984).

Genetic variation at microsatellite loci was estimated per geographical regions (Figure 1) and sub-groups (Figure S1) using the allelic richness (*Ar*), private allelic richness (*pAr*), observed (*H*_*o*_) and expected (*H*_*e*_) heterozygosity. The calculation of *Ar* and *pAr* was performed in *ADZE* (Szpiech et al., 2008) which implements a rarefaction-based standardization procedure to account for differences in sample size between groups. ADZE considers individuals without missing data. Standardized *Ar* and *pAr* were thus computed for a sample size of 18 gene copies for geographical regions and sub-groups. Geographical sub-groups with less than 10 individuals (i.e. 18 gene copies) were discarded. *H*_*o*_ and *H*_*e*_ were first computed with *diveRsity* without applying any rarefaction standardization. Differences in genetic diversity between geographical regions were tested using a Wilcoxon signed-rank test for paired samples. Bonferroni corrections were applied to adjust significance levels for multiple tests. For the mtDNA genetic variation, we estimated nucleotide diversity (*π*), Watterson’s *θ*_*W*_, and haplotype diversity (*Hd*) using *DnaSP*.

While the estimations of *Ar* and *pAr* with *ADZE* already implement a standardization to account for differences in sample size among groups, calculations for the other statistics did not. Differences in sample size can significantly impact the values of genetic diversity estimators (Goodall-Copestake et al., 2012). Therefore, we also applied a rarefaction procedure (Sanders, 1968) using a custom R script to calculate *H*_*e*_, *F*_*IS*_, *R*_*D*_, *π* and *ϑ*_*W*_ in order to account for differences in sample size among geographic regions. However, while *ADZE* subsamples gene copies for the rarefaction procedure, here we subsampled individuals. For each geographical region and each statistic, we randomly subsampled 14 and 5 individuals 1,000 times–respectively for the statistics relative to microsatellite and mtDNA data, respectively. We then estimated the mean and standard error for each statistic and for each geographical region. The rarefaction method was also performed for geographical sub-groups for *H*_*e*_ assuming a standardized sample size of 10 individuals. Sub-groups with less than 10 individuals were excluded (Table S2).

Genetic diversity is expected to decrease with distance away from glacial refugia (Excoffier & Ray, 2008). Therefore, we investigated patterns of spatial variation in various genetic diversity estimators across the North Atlantic subspecies distribution range. We plotted the mean standardized values for *H*_*e*_, *Ar, pAr* and *π* on geographical maps using *MARMAP* v1.0.2 (Pante & Simon-Bouhet, 2013). Spatial interpolation between sampling locations was calculated for each genetic diversity index to assess variation across geographical regions and sub-groups using the R script available at http://membres-timc.imag.fr/Olivier.Francois/plot.admixture.r.

### 2.5 Isolation by distance

Spatially restricted dispersal across generations is expected to promote genetic differentiation among individuals and could create a pattern of isolation-by-distance (IBD) (Rousset, 1997). IBD describes a pattern of genetic differentiation between individuals or populations increasing with geographic distance. In other words, under IBD, populations living closer to each other are expected to be genetically more similar than populations farther away. Such IBD was detected previously in harbor porpoises on each side of the North Atlantic (Fontaine et al., 2007; Rosel et al., 1999a) and is thus expected to occur at the scale of the entire North Atlantic. Furthermore, sex-biased dispersal is widely documented in cetacean species with males generally dispersing more than females (e.g., Dall’s porpoises) (Escorza-Treviño & Dizon, 2000). It has been suggested for harbor porpoises in the Northwest (Rosel et al., 1999a) and Northeast Atlantic (Andersen et al., 1997; Andersen et al., 2001; Tiedemann et al., 1996; Walton, 1997). Sex-biased dispersal can strongly influence population genetic structure (Prugnolle & de Meeus, 2002). Hence, if females disperse less than males across generations, for example due to female philopatry, IBD is expected to be stronger in females than in males, and stronger in mtDNA than in microsatellite loci (Prugnolle & de Meeus, 2002). We assessed the occurrence of IBD by testing the correlation between estimates of genetic differentiation between populations with geographic distance and compared IBD patterns between maternally inherited (mtDNA) and biparental inherited (microsatellites) markers. IBD was tested at the scale of geographical regions and sub-groups. Sub-groups containing less than 10 individuals were not considered (Table S2).

Genetic differentiation at microsatellite loci between populations was estimated as *F*_*ST*_/(1-*F*_*ST*_) which is expected to increase linearly with increasing geographical distance under IBD (Rousset, 1997). We used Weir and Cockerham’s *F*_*ST*_ for microsatellites and *Φ*_*ST*_ for mtDNA. We computed marine geographical distances that account for the shortest path by sea, because a Euclidian distance would poorly describe the actual geographical distance separating pairs of sampled locations in the marine environment. We used *fossil* v.0.3.7 (Vavrek, 2011) and *MARMAP* to estimate these marine geographic distances. Harbor porpoises are usually found on the continental shelf between -10m and -200m, but they have been observed also in high abundance up to -650m in some regions (Skov et al., 2003). Therefore, we constrained the computation of marine distances between -10m and -650m in order to eliminate the possibility of movement across deep basins (Figure S2).

To visualize potential fracture in the spatial distribution of the genetic distance between populations, we plotted genetic distance against marine geographic distances for all population pairs. Then, we focused only on North Atlantic populations (*P. p. phocoena*) to test specifically for IBD, excluding individuals south of the northern Bay of Biscay to avoid the effect of inter-subspecies comparison. We used a Mantel test to assess the correlation significance between genetic and geographic marine distances using 1×10^6^ permutations in the R package *ade4* v1.7 (Dray & Dufour, 2007). We also assessed the relationship between average relatedness (between pairwise sub-groups) and marine genetic distances using the same approach. Average relatedness was assessed using the microsatellite data and the Wang’s estimator of the coefficient of relatedness (Wang, 2002) in the R package *related* v.1.0 (Pew et al., 2015).

### 2.6 Connectivity and gene flow across North Atlantic populations

We further quantify the amount of dispersal, connectivity and contemporary gene flow between geographical regions and sub-groups that included at least 10 individuals using the ‘*divMigrate*()’ function of the R package *diveRsity*. This method relies on the detection of the direction of genetic differentiation, here the *G*_*ST*_ statistics (Nei, 1973), to infer the asymmetric pattern of migration rate (*m*) between groups (Sundqvist et al., 2016). We tested whether gene flow was significantly asymmetric between groups using 5,000 bootstrapped genotype resampling. The results were visualized as a network drawn using *igraph* v.1.2.4.1 (Csardi & Nepusz, 2006) and *popgraph* v.1.5.1 (Dyer, 2017) in R. Nodes in the network represent populations and edges indicate migration rates from one population to another. Networks were constructed at the percolation threshold, *i*.*e*., the highest distance until the network collapses (Rozenfeld et al., 2008). Contemporary effective population sizes (*N*_*e*_) in each geographical region were estimated with *NeEstimator* v2.1.3 (Do et al., 2014) based on LD between microsatellites loci (Waples & Do, 2010). As recommended by the authors, alleles with a frequency lower than 0.02 were filtered out (Waples & Do, 2010). The contemporary effective number of migrants per generation (*2*.*Ne*.*m*) between the geographical regions and the sub-groups was estimated by combining *N*_*e*_ and *m* estimates.

### 2.7 Population demographic changes

In order to assess genetic evidence for effective population size (*N*_*e*_) changes based on mtDNA data, we estimated Tajima’s *D* and Fu and Li’s *D\** in each geographical region using *DnaSP*. These two statistics assess the deviation from the site frequency spectrum from the pattern expected under a neutral constant-size model. For microsatellite loci, evidence of *N*_*e*_ changes in each geographical region was assessed using the Garza and Williamson ratio (*M*_*GW*_) (Garza & Williamson, 2001), which compares the number of alleles to the range in allele size to detect evidence of population contraction. The per region *M*_*GW*_ value was estimated using the *Mratio*() function in *R* available at https://rdrr.io/github/romunov/zvau/man/Mratio.html. *M*_*GW*_, *D* and *D\** were calculated for all the samples and for each geographic region using the same rarefaction approach as described above estimate genetic diversity while accounting for sample size heterogeneity among groups.

### 2.8 Predictive suitable habitat modelling

We used the AquaMaps species distribution modelling approach (Kaschner et al., 2011; Ready et al., 2010) in order to reconstruct the suitable habitat for harbor porpoises at three time periods: at present, during the LGM using on the GLAMAP project data (Schäfer-Neth & Paul, 2004) and by the year 2050, under the most aggressive scenario (Representative Concentration Pathways, RCP8.5) for global climate models of the Intergovernmental Panel on Climate Change (Schwalm et al., 2020). AquaMaps is a bioclimatic model that combines occurrence data (e.g., visual observations, stranding records) with available expert knowledge on species preference and tolerance to different environmental parameters and generates predictions of the probability of occurrence for a target marine species. The preferred habitat can be estimated based on a predefined set of environmental parameters including water depth, sea surface temperature, salinity, primary production, sea ice concentration, and proximity to land. We used mean annual average values for the parameters for the three periods. This was subsequently projected into geographic space as relative probability of occurrence in a global spatial grid of 0.5+ mesh size. The projected predictions of the relative environmental suitability for harbor porpoises into geographic space links habitat preferences to local conditions using environmental data for different time periods and assumes no changes in species-specific habitat usage over time.

In this study, we used a slightly modified version of the original AquaMaps model (available at www.aquamaps.org; Kaschner et al. 2019). Specifically, primary production was excluded from the model, as there are no corresponding data for this parameter for the Pleistocene. AquaMaps has been previously used to hindcast suitable habitat predictions during the LGM for various cetacean species such as killer whales (Morin et al., 2015), common bottlenose dolphins (Nykänen et al., 2019), and narwhals *Monodon monoceros* (Louis et al., 2020).

We included sea ice concentration, depth, and sea surface temperature in the final envelopes as these parameters are known to drive the distribution of cetaceans (Kaschner et al., 2006) (see also the model including salinity in Figure S16 and S17). Using the environmental envelopes, we computed predictions of habitat suitability focusing only on the North Atlantic and adjacent seas for the distribution of harbor porpoises and generated maps using *ggplot*. We calculated and compared the mean latitude of suitable habitat between the three periods using Mann-Whitney U tests. Furthermore, we calculated the size of the suitable habitat for each time period.

## 3. Results

### 3.1 Mitochondrial phylogeography

The mtDNA alignment of 148 sequences (excluding the outgroup) contained 359 segregating sites, including 118 singletons and 241 parsimony informative sites defining 110 distinct haplotypes (Table S3 and S4). Among 88 nucleotide substitution models tested with *jModelTest* v2.1.10, the *HKY+G* substitution model (*G*=0.17) was identified as best fitting the data according to the *BIC* criterion. The maximum-likelihood tree confirmed previous reports of how deep genetic divergence was between Pacific and Atlantic porpoises and that no haplotype was shared between the two ocean basins (Figure 2) (Ben Chehida et al., 2020; Rosel et al., 1995). Within the North Atlantic and Black Sea, four main mtDNA lineages equally divergent from each other emerged; three of them corresponded to the previously identified subspecies (Fontaine et al., 2014) (Figure 2): *(i) P. p. relicta* in the Black Sea; *(ii) IBMA* which included two distinct sub-lineages in the upwelling waters of Iberia (IB) and Mauritania (MA); *(iii) P. p. phocoena* composed of the porpoises sampled from the North Sea to the Arctic waters of Norway, and westward across the mid-North Atlantic waters, off Iceland, Western Greenland and then southwards along the North American coasts of Canada, and Gulf of Maine in the US; *(iv)* a fourth not previously reported lineage carried by a single individual from WGLD (Hap 47). We resequenced this WGLD sample twice to ensure this was not an artefact. This lineage was equally divergent from the three other lineages identified in the Atlantic and Black Sea waters (Figure 2). The ancestral nodes of each lineage were highly supported (bootstrap values > 90%; Figure 2). The haplotype network (Figure S3a) and nDMS (Figure S3b) also clearly depicted these four lineages. Net divergence (*d*_*A*_) between this Hap_47 haplotype and the other three lineages ranged from 0.45% to 0.67% (Figure S4a). This level of divergence overlapped with the divergence observed among the other three main lineages of the Atlantic and Black Sea, ranging from 0.43% to 0.67% and is an order of magnitude higher than the divergence observed among sequences within the *P. p. phocoena* (*d*_*A*_ within NAT ≤ 0.04%) lineage. Hap_47 was excluded for downstream analyses of mtDNA genetic diversity because it was clearly not closely related to any of the other NAT individuals and would bias the estimates of genetic diversity.

**Figure 2.**
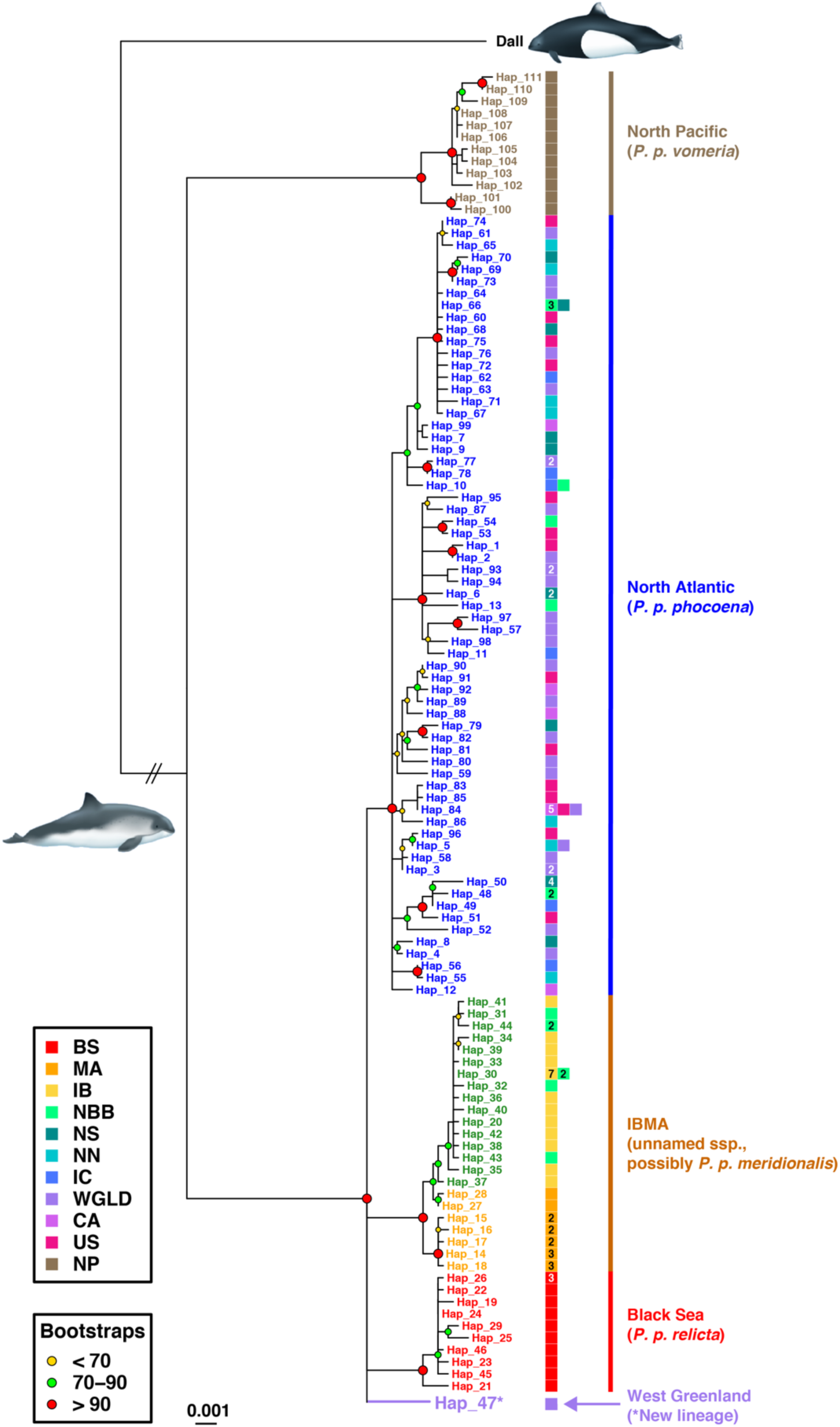
Mitochondrial phylogeny among unique haplotypes estimated using a maximum-likelihood approach. Poorly supported nodes with less than 50% bootstrap support were collapsed. The color-coded labels show the geographic origin of the haplotype. A cryptic lineage (Hap_47) found in West Greenland and distinct from all the others is highlighted in purple. The numbers within the boxes refer to the number of individuals carrying the haplotype. No number means that the haplotype was observed only once. Group acronyms are provided in Figure 1, except for NP: North Pacific.

Aside from Hap_47, we did not observe any clear mtDNA phylogeographic pattern across the North Atlantic within *P. p. phocoena* (Figure 2 and S3). The only remarkable observation was in the southeastern North Atlantic range, in the Bay of Biscay, where haplotypes from *P. p. phocoena* were mixed with those from *IBMA*. This indicates the previously reported hybridization between the two sub-species and the predominantly northward gene flow (Alfonsi et al., 2012; Fontaine et al., 2014) (Figure 2 and S3).

### 3.2 Genetic structure and differentiation

A total of 925 individuals were successfully genotyped at 10 microsatellite loci with less than 7.06% of missing data (Table S1, S2 and S5). Significant linkage disequilibrium (LD) between pairs of microsatellite loci was detected only in 4 out of the 450 tests after applying a Bonferroni correction. These comparisons always included IC. However, the multi-locus LD test did not reveal any significant linkage in any putative groups, with *R*_*D*_ values ranging from -0.006 to 0.06. No significant deviation from Hardy-Weinberg equilibrium (HWE) was observed within the BS, MA, IB and NBB groups with overall fixation index *F*_*IS*_ values ranging from -0.018 to 0.047 (Fisher Exact tests, *p-*value > 0.05; Table S5). In contrast, the NAT group taken as a whole (excluding NBB) showed a significant positive *F*_*IS*_ value (*F*_*IS*_=0.046; *p-*value ≤ 0.01, Table S5), indicating a heterozygosity deficit compared to HWE. This departure from random mating expectation is possibly driven by population structure (i.e., Wahlund effect) (Garnier-Géré & Chikhi, 2001; Wahlund, 1928). When considering each region individually, only IC, NN and NS still exhibited some slight but significant departures from HWE (Table S5).

Consistent with the mtDNA phylogenetic analyses and previous works (Fontaine et al., 2014), the distinct mtDNA lineages of harbor porpoise identified in the North Atlantic and Black Sea waters also formed clearly distinct genetic clusters when analyzing microsatellite variation using multivariate and Bayesian clustering analyses (Figure 3, S5 to S7): (i) *P. p. relicta*; (ii) *IBMA* including the two sub-groups IB and MA and (iii) *P. p. phocoena*. The first principal axis of variation or discrimination (PC1 in Figure S5a, sPC1 in Figure 3a and DF1 in Figure 6c,f) split *P. p. relicta* in the Black Sea from the rest of the North Atlantic porpoises. The second axis (PC2 in Figure S5ab, sPC2 in Figure 3a and DF2 in Figure S6c,d,f,g) discriminated *IBMA* from *P. p. phocoena*. Each multivariate method placed the individuals from the NBB region at an intermediate position between the *IBMA* and *P. p. phocoena* lineages, consistent with their admixed genetic background previously reported (Fontaine et al., 2014; 2017). The global sPCA test assessing the presence of genetic clines or clusters also confirmed that the two first positive sPCs (Figure 3a) were significant (*p-*value ≤ 0.002). By contrast, none of the negative sPCs were significant in the local sPCA test (*p-*value ≥ 0.867, Figure 3a). Focusing only on *P. p. phocoena* (NAT) individuals, none of the sPC were significant (*p-*value > 0.300). Likewise, the DAPC did not reveal any evidence of population subdivision within the *P. p. phocoena* (NAT) subspecies (Figure S8). Also, the distinction between MA and IB, clearly apparent at the mtDNA level (Figure 2 and S3), was also visible in the DAPC using DF3 (Figure S6d,g) and sPC3 (result not shown).

**Figure 3.**
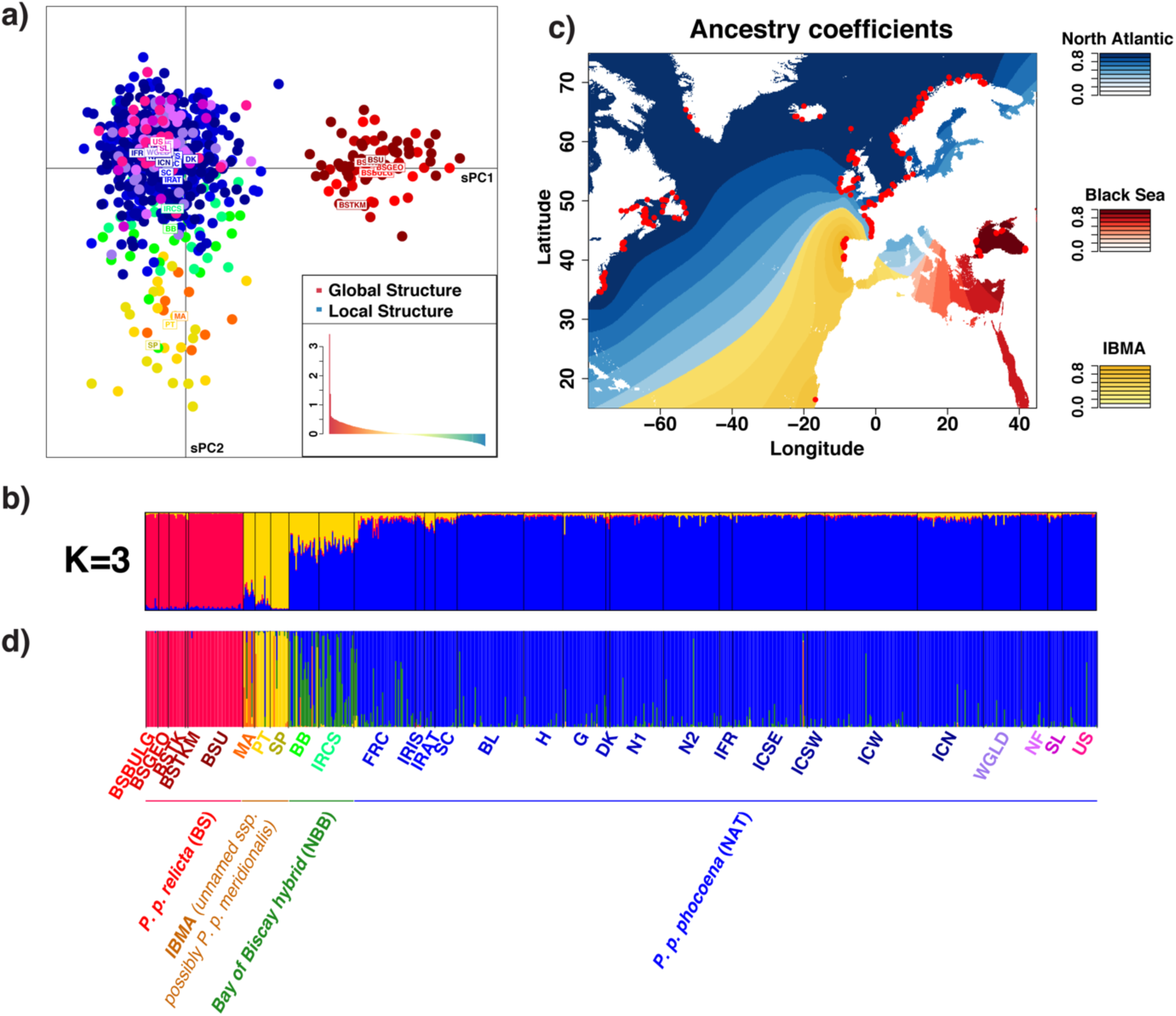
Genetic structure of harbor porpoises in the North Atlantic and Black Sea based on 10 nuclear microsatellite loci. (a) Scatter plot showing the first two spatial principal components (sPCs) of a spatial principal component analysis (sPCA). The inset corresponds to the positive and negative eigenvalues of the sPCA which depicts the global and local genetic structure, respectively. (b) Barplot showing the individual genetic ancestry proportions to each cluster estimated from the *STRUCTURE* analysis at K=3 excluding individuals with missing data. (c) Interpolated map of the genetic ancestry coefficients inferred from the clustering analysis of *STRUCTURE* at K=3. (d) Barplot showing the DAPC cluster membership probability excluding individuals with missing data. BSBULG= Black Sea Bulgaria. BSGEO= Black Sea Georgia. BSTKM=Black Sea Turkey Marmara Sea. BSTK=Black Sea Turkey. BSU=Black Sea Ukraine. MA=Mauritania. PT= Portugal. SP= Spain. BB=Bay of Biscay. IRCS= Celtic Sea. FRC=France Channel. IRIS=. Irish Sea. IRAT=Irish Atlantic. SC=Scotland. BL=Belgium. H=Holland. G=Germany. DK=Denmark. IFR=Faroe Island. N1=Norway South. N2=Norway North. ICN=Iceland North. ICSE=Iceland South East. ICSW=Iceland South West. ICW=Iceland West. WGLD=West Greenland. NF= Newfoundland. SL=Saint Lawrence. US=United States.

The Bayesian clustering analysis of *STRUCTURE* (Figure 3b-c and S7a,c) provided consistent results over 10 replicated runs performed for each number of cluster (*K*) tested and was consistent with the multivariate analyses. The probability of the data greatly increased until K=3, which showed the highest values on average (Figure S7b). At K=2, the analysis split *P. p. relicta* from the remaining porpoises and at K=3, *IBMA* split from *P. p. phocoena* (Figure S7a). Beyond K=3, no further subdivision was observed. Fontaine et al. (2014) showed that the two groups of *IBMA* in MA and IB were more closely related to each other than with the other porpoises and could be discriminated from each other only when analyzing them separately from the other porpoises. Also similar to previous studies (Fontaine et al., 2014) and the multivariate analyses, NBB showed an admixed genetic ancestry with an equal contribution from *IBMA* and *P. p. phocena* subspecies (Figure 3b and S7a). No finer subdivision was observed over the whole area covered by *P. p. phocoena*, even when running *STRUCTURE* focusing only on *P. p. phocoena* individuals (Figure S7c). The most likely number of ancestral genetic clusters within *P. p. phocoena* was one, as suggested by the likelihood of the data at K=1 (Figure S7d).

Interestingly, the individual carrying the fourth major mtDNA lineage (Hap_47) in Western Greenland (WGLD) could not be distinguished from other NAT porpoises based on the microsatellite data. This may just be due to the fact that only one individual of a distinct cluster has been sampled or this individual may also share most of its genetic ancestry with the NAT group at the nuclear loci, but not at the mtDNA locus.

Genetic differences in microsatellite allelic frequencies (Figure 4a, S9 and S10) and mtDNA haplotype frequencies (Figure 4b, S4b,c and S10) between subspecies were all highly significant (*p-* value < 0.001) and much higher than the comparisons among demes within each subspecies (Figure 4a,b, S9, and S10). Among subspecies, *F*_*ST*_ values for microsatellite loci ranged between 0.09 and 0.30. For the mtDNA, *ɸ*_*ST*_ values ranged from 0.61 to 0.90. The highest *F*_*ST*_ values were observed between BS and *IBMA* and lowest between IB and MA. Within *P. p. phocoena* (NAT) subspecies, microsatellite *F*_*ST*_ values among the geographical regions (Figure S9a) and sub-groups (Figure S9b) were much smaller and none of the pairwise comparisons were significantly different from zero. In contrast, some demes from NWA and NEA were significantly differentiated for the mtDNA *F*_*ST*_ and *Φ*_*ST*_ comparisons (Figure S4b,c).

**Figure 4.**
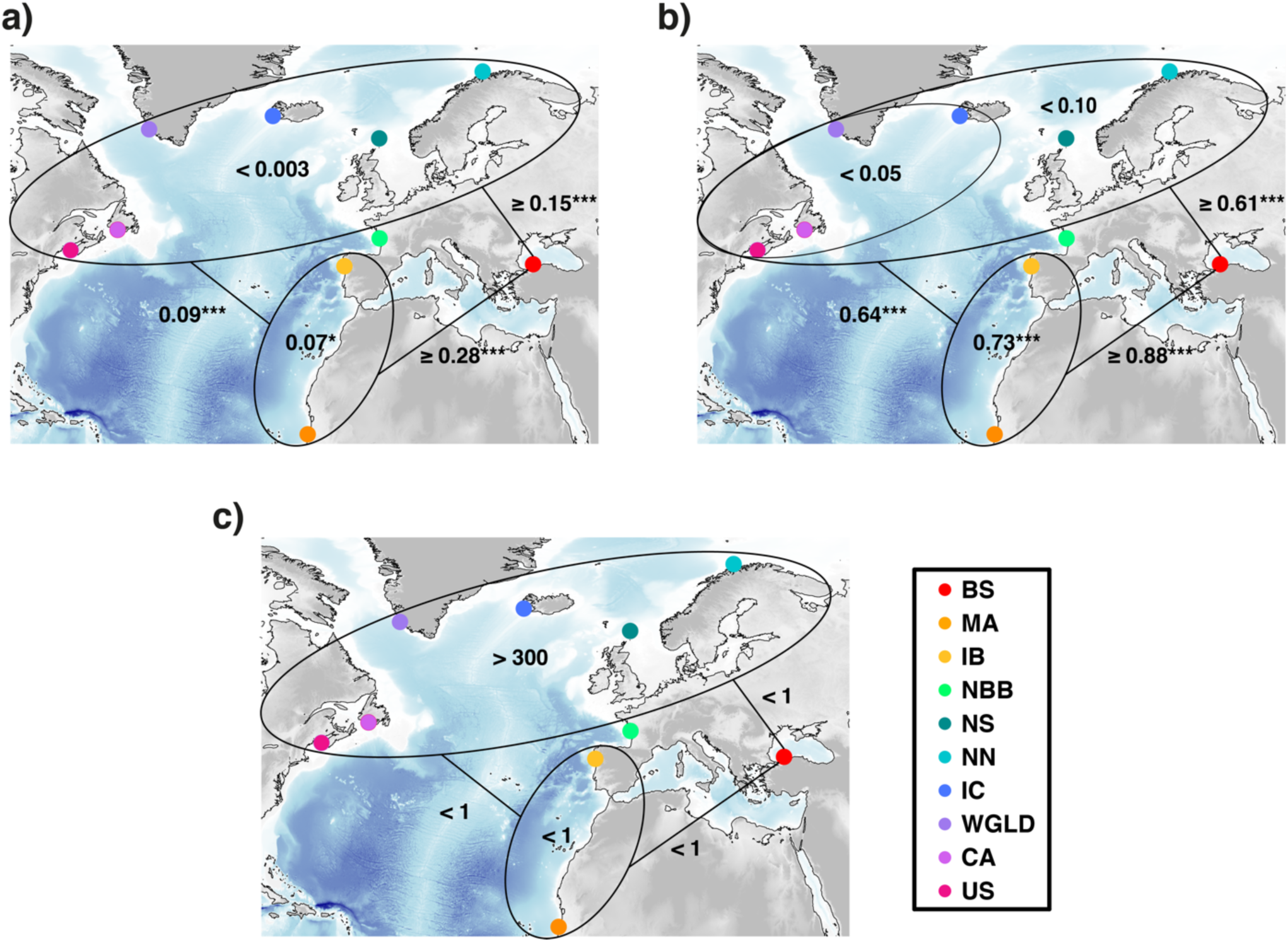
Genetic differentiation at microsatellite (a) and mtDNA (b) loci, expressed as the average pairwise *F*_*ST*_ and *ɸ*_*ST*_ values, respectively. (c) Connectivity among regional groups of harbor porpoises is displayed as the average effective number of migrants (*2*.*Ne*.*M*) per generation. *= *p-value* ≤ 0.05. ***= *p-value* ≤ 0.001. The acronyms are provided in Figure 1.

Visualizing the genetic differences as a function of the marine geographic distance showed that the increase in genetic differentiation between *P. p. phocoena* (NAT), *IBMA* or *P*.*p. relicta* (BS) subspecies is not simply an effect of the increasing geographic distance (Figure S10). The relationship between genetic and geographic distance revealed large jumps in *F*_*ST*_ or *ɸ*_*ST*_ values, reflecting genetic discontinuities indicative of barriers to gene flow or secondary contact between subspecies as previously noted by Fontaine et al. (2007; 2014; 2017). We did not detect any such discontinuity between demes within *P. p. phocoena*. The intermediate position of NBB was again clearly illustrated with lower levels of genetic differentiation than the among subspecies comparisons but higher than the within cluster comparisons (Figure S10). This variation in genetic differentiation among demes between and within subspecies translates variation in gene flow between them.

### 3.3 Population connectivity

We quantified gene flow using microsatellite markers by estimating local effective population sizes (*N*_*e*_, Table S6) and migration rate (*m*, Table S7) to derive the effective number of migrants per generation (2.*Ne*.*m*, Table S8). Consistent with previous studies (Fontaine et al., 2014; 2012; 2010), estimates of *N*_*e*_ (Table S6) were lowest in the two *IBMA* populations (IB and MA with *Ne* < 60 individuals), low in the hybrid NBB (*N*_*e*_*=*217), intermediate in *P. p. relicta* (*N*_*e*_*=*504), and larger in *P. p. phocoena* (*N*_*e*_ > 800 individuals). Furthermore, *N*_*e*_ differed substantially among demes within *P. p. phocoena*. Demes in central North Atlantic (IC/WGLD < 925 individuals) displayed slightly smaller *N*_*e*_ than in the Eastern (NS/NN > 1500 individuals) and Western (US/CA > 1482 individuals) waters.

Estimates of migration rates (*m*, Table S7 and Figure S11) and effective number of migrants per generation (Table S8 and Figure 4c) showed no evidence of gene flow between *P. p. relicta* and the other subspecies in the Atlantic (*IBMA* and *P. p. phocoena, m* < 0.001 and 2.*Ne*.*m* < 1 individual per generation). Likewise, the two lineages IB and MA of *IBMA* in the upwelling waters showed very limited amount of gene flow between them (*m* ≤ 0.021, Table S7 and Figure S11; *2*.*Ne*.*m* < 1 individuals, Table S8 and Figure 4c). Asymmetric gene flow was detected between *IBMA* and *P. p. phocoena*, flowing especially from IB to *P. p. phocoena* demes with NBB serving as a hub in the population network (Table S7 and Figure S11). Estimated migration rates from *IBMA* and *P. p. phocoena* were several times larger than in the reverse direction (Table S7). However, due to the low *N*_*e*_ in *IBMA* lineages (Table S6), the effective number of migrants remained low (2.*Ne*.*m* < 1 individuals) in both directions. Gene exchanges from NBB to *P. p. phocoena* ranged from 0.083 to 0.174 and were larger than those from NBB to *IBMA* (from 0.021 to 0.064). Finally, although not statistically significant, *m* values from NBB to *IBMA* were two to three times lower than the converse. Among *P. p. phocoena* (NAT) demes, we detected a clear signal of migration with *m* varying from 0.18 to 0.9 (Table S7) and 2.*Ne*.*m* exceeding 300 individuals per generation across the whole North Atlantic (Table S8). We observed no difference between the different sectors (West, center and East) of the North Atlantic (Table S7 and S8).

### 3.4 Isolation by distance

Despite the high connectivity among *P. p. phocoena* demes, we tested whether individual dispersal across generations was geographically restricted and could generate an IBD pattern at mtDNA and microsatellite loci at the scale of the North Atlantic. This could explain the departure from random mating expectations detected at microsatellite loci with the significant *F*_*IS*_ value when considering *P. p. phocoena* as a whole (Table S5). We detected a strong IBD at the mtDNA marker characterized by an important increase in genetic differentiation with geographic distance among the *P. p. phocoena* geographical groups (*r*^2^ = 0.68; *p-*value ≤ 0.001; Figure 5a). In contrast, a much weaker IBD was detected at the nuclear microsatellite loci among *P. p. phocoena* demes, and the pattern was only significantly detectable when considering the finest level of geographic subdivision (*r*^*2*^ = 0.04; *p-*value ≤ 0.004; Figure 5b), not with the geographical regional subdivisions (*p-*value > 0.05; Figure S12). Consistent with this weak but significant IBD pattern at the microsatellite level, we also detected a negative relationship in relatedness as geographical distance increases between pairs of *P. p. phocoena* demes (*r*^*2*^=0.05; *p-*value ≤ 0.03; Figure 5c). The IBD patterns highlighted in this study suggest that *P. p. phocoena* is not a panmictic lineage because of restricted intergenerational dispersal over its entire geographical range.

**Figure 5.**
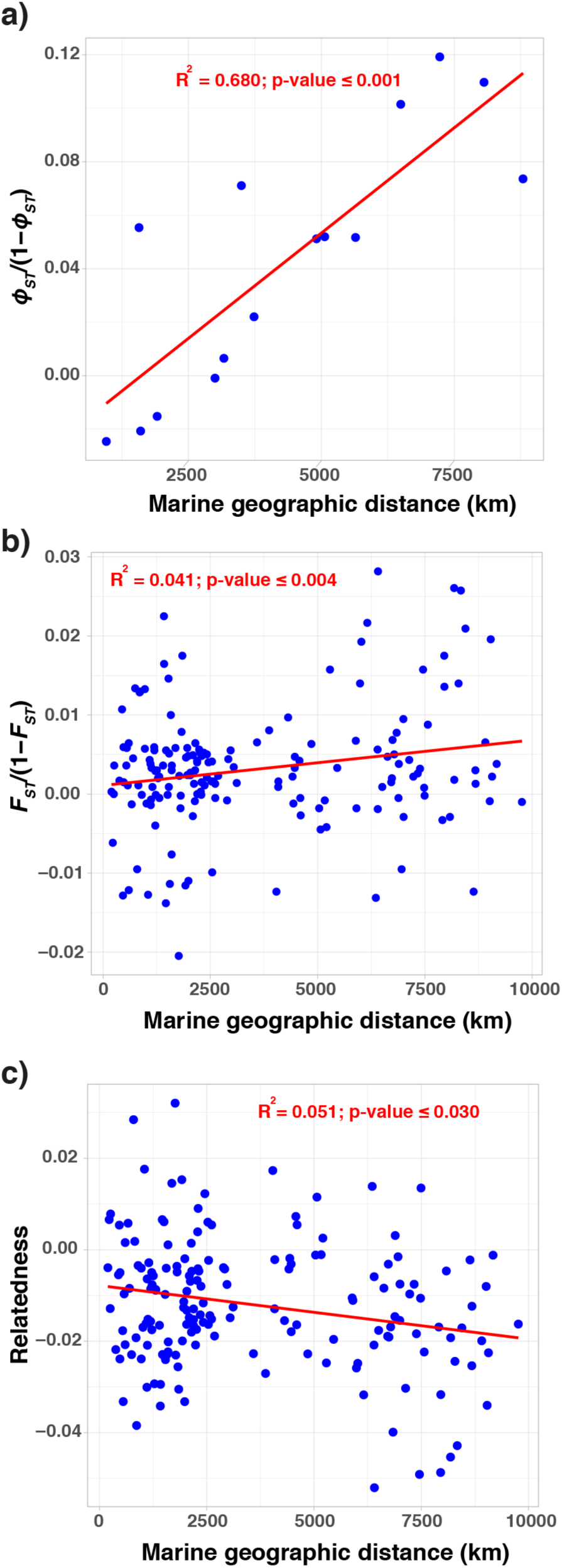
Pattern of isolation by distance among subpopulations of *P. p. phocoena* in the North Atlantic. Relationship between the unbounded estimator of genetic differentiation among geographical regions or sub-regions and their marine geographic distances for (a) mtDNA and (b) microsatellite loci. (c) Relationship between the Wang’s relatedness estimator (Wang, 2002) and the marine distances among sub-groups. Red lines show the regression lines and R^2^ provide the determination coefficient.

### 3.5 Spatial variation in genetic diversity and demographic changes

We investigated spatial variation in genetic diversity in order to assess the evidence for potential glacial refugia in *P. p. phocoena* together with a potential leading-edge effect that could have accompanied the post-LGM recolonization of the Nordic waters. MtDNA genetic diversity (Table S4, Figure S13a and S14a), quantified using the haplotype diversity (*Hd*) and two estimators of nucleotide diversity (*π* and *θ*_*w*_), was lowest in the two populations of *IBMA*, slightly higher in *P. p. relicta*, and highest in *P. p. phocoena*. At the contact zone between *IBMA* and *P. p. phocoena*, NBB displayed the highest mtDNA diversity of all geographical regions, which is expected given the mixture of divergent haplotypes from the two sub-species in that region. North of the Bay of Biscay, the genetic diversity was more homogeneous across the North Atlantic among *P. p. phocoena* demes. It was slightly higher in NS, IC and WGLD. However, when accounting for difference in sample sizes, standard errors of all estimates overlapped (Table S4, Figure S13a and S14a) suggesting no significant differences in mtDNA diversity among *P. p. phocoena* demes.

The level of microsatellite diversity was comparable between *IBMA* and *P. p. relicta* (Table S2 and S5, Figure S13b, S14b-d, S15). This is suggested by the lack of significant differences in allelic richness (*Ar*), private allelic richness (*pAr*), and expected heterozygosity (*He*) (Wilcoxon Signed Rank tests, WSR, *p-*value > 0.05 for all pairwise comparisons). These sub-species displayed significantly lower diversity than the admixed porpoises (NBB) and those from *P. p. phocoena* (WSR, *p-*value ≤ 0.002 for the three diversity measures). NBB porpoises showed lower *Ar* and *He* than *P. p. phocoena* (WSR, *p-*value ≤ 0.03) and lower *pAr* than US, NS and CA, similar to IC and NN and higher than WGLD (WSR, *p-*value ≤ 0.009). While *Ar* and *He* statistics were comparable among *P. p. phocoena* demes, *pAr* was significantly higher in US (WSR, *p-*value ≤ 10^−5^) and lower in WGLD (WSR, *p-*value ≤ 0.001). NS also displayed significantly higher *pAr* than WGLD, IC and NN (WSR, *p-*value ≤ 0.001). The remaining geographical regions showed no significant difference in *pAr* (WSR, *p-*value > 0.05).

We detected significant evidence of change in effective population size in *P. p. relicta* (BS) and IB population within *IBMA* as shown by significant negative values for both Tajima’s *D* and Fu & Li’s *D\** statistics (Table S4, Figure S13a). MA did not display any significant departure from neutral expectations, and neither did any region within the *P. p. phocoena* range (Table S4, Figure S13a) for Tajima’s *D*. WGLD and US showed a significant negative *D\** when considering all the samples, but the values did not significantly depart from 0 when using the rarefaction approach (0 being included in the standard error; Figure S13a). Hence, for mtDNA, all *P. p. phocoena* demes seem close to the migration-drift equilibrium suggesting no significant recent changes in demography. At the nuclear microsatellite markers, we found evidence of *Ne* contraction in *IBMA, P. p. relicta* and NBB, with *M*_*GW*_ values statistically smaller than in *P. p. phocoena* (no overlapping standard errors, Table S5 and Figure S13b).

### 3.6 Predictive habitat suitability

Our estimated distribution of core suitable habitats during the present-day reflected well the known distribution of harbor porpoises in the North Atlantic and adjacent seas, especially when considering habitat suitability with a probability larger than 0.3 (Figure 6b; S16 and S17). The suitability envelope ≥0.3 included indeed all the sampling points and all the areas where harbor porpoises have been reported (see the area status report of the NAMMCO & IMR, 2019). Non-zero habitat suitability values were also predicted in the Azorean waters and in the Mediterranean Sea, where porpoises are mostly absent nowadays, but habitat suitability remained mostly under 0.3, except in the Aegean Sea and Alboran Sea where sightings have been occasionally reported (reviewed in Fontaine, 2016; NAMMCO & IMR, 2019 and references herein).

**Figure 6.**
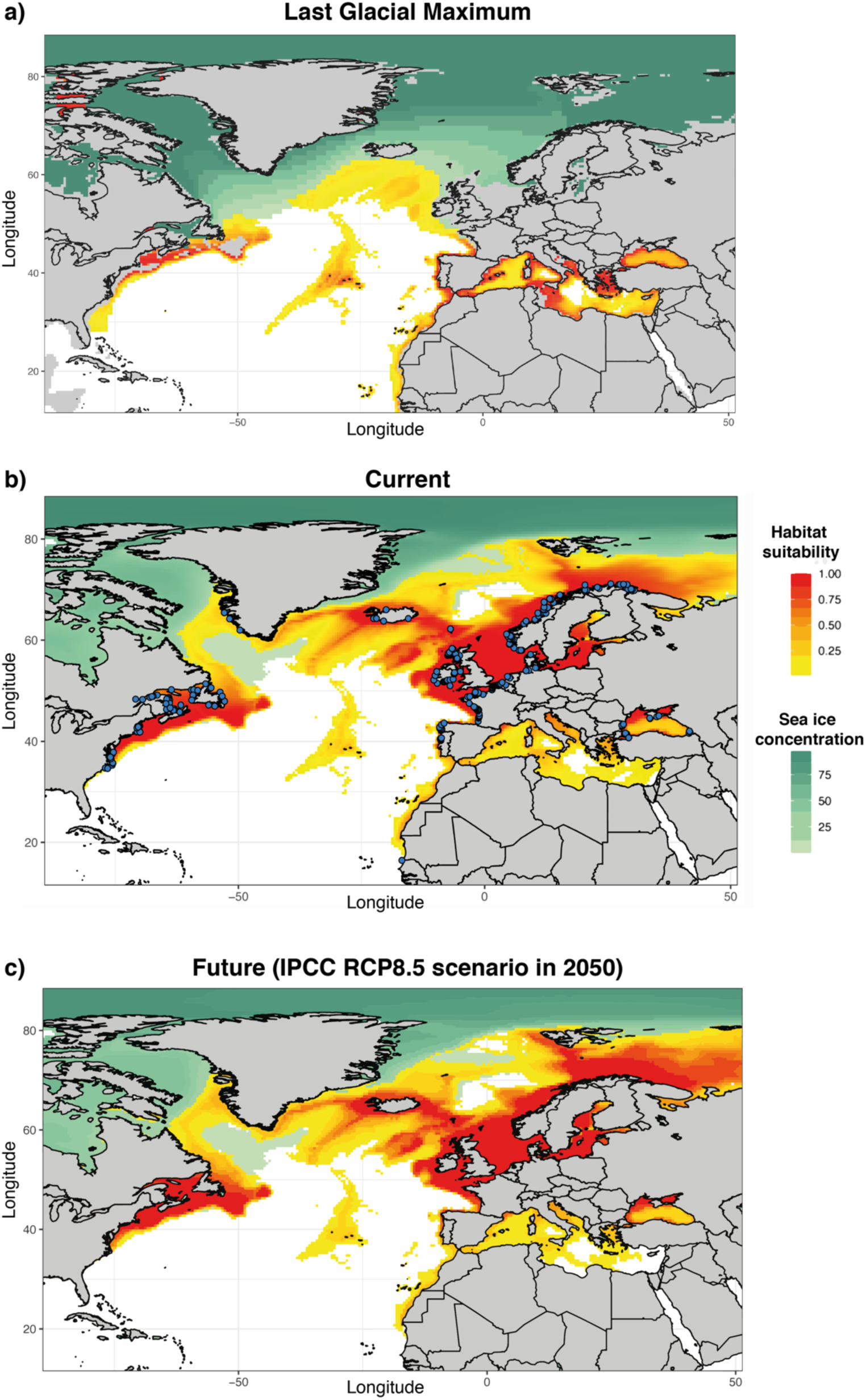
Maps showing the predicted habitat suitability for harbor porpoises throughout the North Atlantic and adjacent seas during three time periods generated using AquaMaps environmental niche modelling and input parameter settings described in Table S9, excluding salinity as predictor. Yellow to red colors represent least to most suitable habitat, respectively, based on the AquaMaps habitat model. Light to dark green colors represent the proportion of sea ice concentrations (%). Emerged lands are displayed in grey. Blue dots on the current map show individual locations of the porpoise samples used in this study.

The AquaMap model predicted that the total surface of the suitable habitats was reduced by a factor of three during the LGM, from ∼15 million km^2^ at present to ∼5 million km^2^ (Figure 6 and S18). The average latitude of habitable zones during the Pleistocene were also significantly compressed and shifted southwards around 40ºN (5^th^ and 95th percentiles: [30ºN - 48ºN]), compared to the significantly higher and broader present-day latitudinal distribution (average at 58ºN, [39ºN -74ºN], *p*-value < 0.001). Noteworthy is the increase in habitat suitability during the LGM (Figure 6a) in the Azorean waters and in the Mediterranean Sea compared to the present-day (Figure 6b). In contrast, a significant part of present-day habitats was unavailable during the LGM. These include waters off Greenland, Iceland, Norway, North Sea, Baltic Sea, but also the Black Sea which was only reconnected back to the Mediterranean Sea ∼7000 yrBP (Fontaine et al., 2012; 2010; reviewed in Fontaine 2016).

The model did not reveal any significant difference between present and future distributions of suitable habitats (Figure 6 and S18). In contrast, habitat size appears even to slightly increase for the year 2050 (Figure S18).

## 4 Discussion (3688)

Quaternary glaciations deeply impacted the genetic structure of terrestrial and marine organisms (Hewitt, 2000), including cetacean species like harbor porpoises in the North Atlantic (Fontaine, 2016). The split between the two subspecies on each side of the Mediterranean Sea during the LGM was one of the most dramatic events in the species evolutionary history, leaving one relict subspecies in the Black Sea (*P. p. relicta*) and another one (*IBMA*) in the upwelling waters of Iberia and Mauritania (Fontaine, 2016; Fontaine et al., 2014). The third subspecies, *P. p. phocoena*, is the most widely distributed in the coastal waters of the North Atlantic, north of the Bay of Biscay, but its population structure and evolutionary history remained contentious despite numerous genetic studies (Andersen, 2003; Andersen et al., 2001; Fontaine et al., 2007; 2014; 2017; Rosel et al., 1995; 1999a; 1999b; Tolley & Rosel, 2006; Tolley et al., 2001; Wang et al., 1996). Limited sample size, disparate geographic sampling and genetic markers, lack of resolution in genetic markers, and heterogeneous methodologies (reviewed in Andersen, 2003; Fontaine, 2016; NAMMCO & IMR, 2019) contributed to maintaining an incomplete understanding of the population structure and evolutionary history across the North Atlantic. The present study aims at filling this gap by compiling a fairly comprehensive sampling over the entire species distribution range in the North Atlantic, focusing on a synchronous cohort collected between 1990-2000 and including a spatio-temporal perspective of how suitable habitats changed since the LGM and how it will evolve in the near future under the most aggressive (RCP8.5) scenario of the IPCC.

### 4.1 A new mtDNA lineage in harbor porpoises from Western Greenland waters

The discovery of a new divergent mtDNA lineage not previously uncovered in the North Atlantic is a major finding of this study. This haplotype, carried by a single individual from western Greenland (WGLD), displayed the same level of divergence as the three other subspecies previously identified in the North Atlantic and Black Sea (Figure 2, S3a,b and S4a). The phylogenetic tree (Figure 2) ruled out the possibility that this haplotype could come from migrants from the North Pacific. The genetic divergence between Pacific and Atlantic porpoises is indeed far larger than the divergence observed between lineages present in the North Atlantic and Black Sea waters. Divergence time estimates between porpoises from the two ocean basins was consistent with a split between 0.7-0.9 Myr ago (Tolley & Rosel, 2006). This contrasts with the polytomy observed among the four Atlantic-Black Sea mtDNA lineages in the phylogenetic tree suggesting a rapid split between the subspecies previously estimated during the height of the LGM ∼20 kyr BP (Fontaine, 2016; Fontaine et al., 2014; 2010).

This new mtDNA lineage in Western Greenland is reminiscent of the recent discovery of a cryptic ecological group of harbor porpoises in the exact same area by Nielsen et al. (2018). Satellite tracking of 30 WGLD individuals revealed that they were using oceanic habitats and were diving to depths that enable mesopelagic foraging, contrasting with the demersal feeding habits in shallow waters (within 200m) usually reported so far for this species. These distinctive migratory and diving behaviors suggested that WGLD porpoises could belong to a unique oceanic ecotype, distinct from neighboring *P. p. phocoena* populations. Moreover, Nielsen et al. (*in prep*.) found, based on preliminary analyses of single nucleotide polymorphism (SNP) markers, that individuals from this ecotype displayed shallow but significant genetic differentiation and admixture with the neighboring populations. The highly divergent haplotype uncovered here in one individual sampled opportunistically between 1990 and 2000 may belong to this ecologically distinct group. This particular individual was not distinguishable with the nuclear microsatellite loci from other *P. p. phocoena* demes. The shallow nuclear divergence seems thus in contrast with the relatively deep mtDNA divergence. This likely comes from the fact that the mtDNA molecule does not recombine, in contrast to nuclear markers. MtDNA traces maternal ancestry back to the Most Recent Common Ancestor (MRCA) without being broken down by recombination and thus without being impacted by homogenizing effects of gene flow with other lineages. Future studies with a larger sample size shall examine in greater detail the genetic structure and evolutionary history of this peculiar group.

The equal degree of divergence between this new mtDNA lineage and the other Atlantic and Black Sea lineages (Fig 2, S3a,b and S4a) suggests that they all split almost at the same time during the LGM (Fontaine et al., 2014). They must thus have remained isolated in allopatry long enough to allow mtDNA divergence to build up. We may speculate that this cryptic lineage could have emerged during the LGM in the oceanic waters surrounding the Azores, where the offshore behavior of this group could have evolved. Even if once considered highly improbable, the occurrence of an offshore porpoise population in the Azores has been suggested multiple times in the past, with even one stranding case reported in January 2004 on the Azorean coasts (Barreiros et al., 2006). Our suitable habitat modelling showed that Azorean waters were also suitable for porpoises during the LGM (Figure 6; S16 and S17). Interestingly, the Azores were a glacial refugia for multiple benthic and pelagic marine species in the North Atlantic (Maggs et al., 2008), such as the thornback rays (*Raja clavata* (L.), Rajidae) (Chevolot et al., 2006), Montagu’s blenny (*Coryphoblennius galerita*, Blenniidae) (Francisco et al., 2014) or ballan wrasse (*Labrus bergylta*) (Almada et al., 2017). One could thus speculate that the fourth mtDNA lineage of the oceanic harbor porpoises in WGLD waters could come from a glacial refugium in the Azores.

### 4.2 No genetic evidence of leading-edge effect in P. p. phocoena

It was originally hypothesized that harbor porpoises on each side of the North Atlantic could have been isolated from each other in distinct glacial refugia, recolonizing northern waters from different southern refugia (Gaskin, 1984; Rosel et al., 1999b; Yurick & Gaskin, 1987). Our results and previous ones (Fontaine, 2016; Fontaine et al., 2014) suggest that the evolutionary history of the species during the Pleistocene is far more complex in the Atlantic and Black Sea. Porpoises around the Mediterranean Sea, from which descended *IBMA* and *P. p. relicta* (Fontaine, 2016; Fontaine et al., 2014), evolved independently from other Atlantic populations during the LGM, likely driven by environmental changes that deeply impacted habitat suitability in that area (Figure 6, S16 to S18). Furthermore, an oceanic refugium (e.g., in the Azores) is also possible (Figure 6, S16 to S18), and could explain the distinct oceanic ecotype discovered by Nielsen et al. (2018). The evolutionary history of the most widespread *P. p. phocoena* subspecies north of Biscay remained, however, debated. Our results showed that *P. p. phocoena* porpoises on both side of the North Atlantic belong to a same mtDNA lineage with high haplotype diversity. This suggests they formed a coherent genetic group during the LGM. This rules out previous hypotheses that two distinct genetic groups recolonized northern waters with a contact zone somewhere in the middle Atlantic, or between Iceland and Norway (Rosel et al., 1999b; Tolley et al., 2001) or by the Davis Strait (Gaskin, 1984; Yurick & Gaskin, 1987). Here, we show that no such genetic fractures occurred among demes within *P. p. phocoena*, in contrast with the sharp increase in genetic differentiation observed between distinct subspecies, for example in the Bay of Biscay (Figure S10). One could argue that not enough generations elapsed since the LGM for genetic differentiation to capture demographic isolation that could have occurred on each side of the North Atlantic, due for example to incomplete lineage sorting. Such a lag time between demographic and genetic differentiation is known as the “grey zone of population differentiation” and has been investigated using empirical and simulation-based studies (Bailleul et al., 2018; Gagnaire et al., 2015), including cetacean species like harbor porpoises (Ben Chehida et al., 2019). This period of time during which genetics would not capture demographic population differentiation is a function of the local effective population size (*N*_*e*_) that conditions the strength of genetic drift. We showed recently using simulations that such a “grey zone” would be short in cetacean species like porpoises, displaying relatively low fecundity and small *Ne* (Ben Chehida et al., 2019). Assuming that two populations would split with no gene flow between them and that each one has an *Ne* value of 1000 diploid individuals, it would take less than 8 generations (80 years assuming a generation time of 10 years for porpoises) to be able detect highly significant *F*_*ST*_ values using 10 diploid nuclear microsatellite markers and four time less with haploid markers such as the mtDNA. Therefore, the lack of genetic fracture between Eastern and Western North Atlantic demes of *P. p. phocoena* at both mtDNA and nuclear microsatellite markers is not a result of any populations grey zone effect. It rather suggests this subspecies remained unfragmented during the LGM or that any subdivisions that previously existed was erased by dispersal during the post-glacial colonization of northern waters progressively freed from Pleistocene ice.

Another hypothesis previously suggested that porpoises may have retreated and contracted in a southern refugia in the Northwest Atlantic during the LGM, and then rapidly expanded into northern waters, and colonized the Northeast Atlantic and North Sea following the retreat of the last Pleistocene glaciers (Rosel et al., 1999b; Tolley et al., 2001). Such a hypothesis was previously supported by higher genetic diversity at the mtDNA-CR in the Northwest compared to the Northeast Atlantic (Rosel et al., 1999b). Our habitat modelling predictions during the LGM (Figure 6, S16 and S17) showed that habitats remained available in the Northwest Atlantic, but in central and Northeast Atlantic as well. Furthermore, our reassessment of the genetic variation at one quarter of the mitogenome sampled in unrelated individuals and a statistical rarefaction procedure that accounts for difference in sample size among regions did not capture any significant spatial variation in mtDNA genetic diversity nor any significant differences between the Northwest and Northeast Atlantic demes (Table S4, Figure S13a and S14a). Our results suggest instead that no major population contraction occurred in *P. p. phocoena* during the LGM. This agrees with our previous results based on coalescent-based inferences that showed a steady increase in effective population size of *P. p. phocoena* porpoises in the Northeast and Central North Atlantic since the time of the MRCA (T_MRCA_) more than 50kyr ago without any evidence of historical population contractions (Fontaine et al., 2014). This suggests that even if suitable habitats for this subspecies contracted by a factor of three, as suggested by our habitat modelling and shifted southwards (Figure 6, S16 to S18), it did not induce major population contractions or that local loss of diversity was compensated by re-colonization through dispersal among neighboring demes, leaving no detectable imprints on the genetic variation of the mtDNA and microsatellite markers.

Post-glacial re-colonization of the northern habitats released from Pleistocene ice could have generated a leading-edge effect as postulated previously for the North Atlantic harbor porpoises (Rosel et al., 1999b; Tolley et al., 2001). However, our results are not consistent with the expectations under such a model. Under a leading-edge model, populations that have expanded rapidly from a core population are expected to experience loss of genetic diversity due to genetic drift operating through repeated bottleneck / founder events (also called allelic surfing) (Excoffier et al., 2009; Hewitt, 1996, 1999, 2000). Under such a model, genetic diversity decreases progressively from the source populations in the glacial refugium towards populations at the colonization front. We did not observe any evidence of clinal variation in genetic diversity at the mtDNA level in any direction among geographic regions, and especially not from southern Northwest to Northeast Atlantic as would have been expected under a rapid expansion in this direction. Instead, mtDNA diversity was highest in the North Sea, an area among the last to be released from the Pleistocene ice, and lowest in Canadian, US and northern Norwegian coasts (Figure S13a and S14a). Spatial variation in nuclear microsatellite loci was also very comparable among *P. p. phocoena* demes (Figure S13b, S14 and S15). Instead, regional variation in genetic diversity is more consistent with local variation in census population size, with the highest census population sizes reported in the North Sea (NAMMCO & IMR, 2019). This suggests that population genetic variation at both mtDNA and microsatellite loci have reached an equilibrium state between migration and genetic drift, so called migration-drift equilibrium. The highly significant isolation by distance (IBD) that we detected at both mtDNA and nuclear microsatellite loci among *P. p. phocoena* demes provide further evidence that migration-drift equilibrium has been reached in this group (Hutchison & Templeton, 1999). Thus, in contrast with previous studies that suggested a potential leading-edge effect, our results based on relatively geographically extensive sampling show that the genetic structure reflects more a combination of recent intergenerational dispersal and local effective population size rather than a post-glacial expansion wave front from southern habitats.

### 4.3 Isolation by distance, female philopatry and dispersal across the North Atlantic

The re-establishment of migration-drift equilibrium since the LGM could actually be expected given the high dispersal capabilities of the harbor porpoise, in particular when considering the effect across generations. In highly mobile species forming a continuum with no obvious barrier to gene flow, as observed among *P. p. phocoena* demes, high intergenerational dispersal quickly redistributes genetic variation among demes. Leblois et al. (2004) showed using simulations that reliable inference of IBD parameters can be done within a few tens of generations, assuming temporal and spatial fluctuations of demographic parameters have not been too strong nor too recent, which seems to be the case for *P. p. phocoena*. This suggests that migration-drift equilibrium can be restored within that time frame. Assuming a generation time of 10 years for harbor porpoise (Read, 1999), this means that a few hundred years are required to restore migration-drift equilibrium. As a matter of fact, persistence in time of the effect of historical demographic fluctuations strongly depends on various demographic features, but in particular the local effective individual density (*D*) and the variance of intergenerational parent-offspring’s dispersal distance (*σ*^2^). Both parameters combined (*Dσ*^2^) form the neighborhood size (*4πDσ*^2^). This neighborhood size reflects the local effect of genetic drift and gene flow in a stepping-stone population model (Rousset, 2000; Rousset, 1997). The smaller the neighborhood size, the stronger local genetic drift is in the face of intergenerational gene dispersal, and the stronger that IBD becomes. Previously detected in the Northeast Atlantic (Fontaine et al., 2007; Tolley & Rosel, 2006), we showed here that IBD extends over the entire North Atlantic. This clearly shows that individual dispersal across generations is geographically restricted. Therefore, for *P. p. phocoena*, panmixia does not hold at the scale of its complete range. Instead, it represents a continuously distributed system under isolation by distance where panmixia hold only within the neighborhood size, *i*.*e*. the demes.

Interestingly, IBD at the mtDNA locus was ten times stronger than at the nuclear microsatellite loci. Previous studies reported analogous evidence by observing significant genetic differentiation at the mtDNA locus that was weak or absent at microsatellite loci among demes in the Northwest Atlantic (Rosel et al., 1999a) and in the Northeast Atlantic (Andersen et al., 2001; Fontaine et al., 2007; Tolley & Rosel, 2006). All these results indirectly suggest strong female philopatry in porpoises, which would lead to a reduced variance in intergenerational mother-offspring’s dispersal distance (*σ*^2^_mtDNA_) and thus to an increase in IBD strength at the maternally inherited mtDNA locus. In contrast, male-biased dispersal would create highly variable father-offspring’s intergenerational dispersal distances, thus increasing *σ*^2^_microsatellites_. Direct observations of such female restricted dispersal driven by female philopatry in harbor porpoise were recently demonstrated by satellite tracking in West Greenland, the central North Atlantic and in the North Sea (Nielsen et al., 2018). Another often neglected factor explaining the difference in IBD strength between the two types of markers is the difference in effective population size. The maternally inherited haploid mtDNA locus implies a four times smaller effective population size at that locus compared to the biparentally inherited diploid nuclear microsatellite loci. This means that the neighborhood size, and especially the local effective individual density (*D*), estimated for the mtDNA locus is thus also four times smaller than for microsatellite loci. Hence genetic drift is increased in the first compared to the second type of marker. Therefore, the combined effect of reduced intergenerational mother-offspring’s dispersal distance driven by female philopatry and the reduced effective mtDNA population size may explain the dramatic differences observed between the two types of genetic markers.

### 4.4 Conservation and management implications

Harbor porpoises are facing numerous threats driven by human activities (e.g., prey depletion due to overfishing, noise disturbance, pollution, and potentially climate change), but by-catch mortalities in commercial gillnet fisheries is by far the most immediate threat to the species (NAMMCO & IMR, 2019). In some regions, bycatch mortality is believed to be unsustainable. Assessing the incidence is essential to design informative and effective management and conservation plans. The International Whaling Commission (Donovan & Bjørge, 1995; Gaskin, 1984) and other studies (Evans & Teilmann, 2009; Fontaine, 2016; NAMMCO & IMR, 2019; Rosel et al., 1999a; Rosel et al., 1999b) have suggested that local demes or subpopulations across the North Atlantic should be treated as distinct assessment or management units (MUs). The delimitation of MUs was originally made based on oceanographic, ecological, and practical considerations. It was later backed-up by multiple lines of evidence ranging from genetic differences to differences in skull morphometrics, life history parameters, stable isotope and fatty acid signatures, and movements revealed from radio-tracking (Andersen, 2003; Evans & Teilmann, 2009; Lockyer, 2003).

Previous definitions of MUs proposed by the IWCs and ASCOBANS were recently reviewed by the IMR & NAMMCO report (2019) and still hold for most part in the face of the present results. The status of the Iberian and Mauritanian populations, the *IBMA* porpoises, was already underlined previously as ecologically distinct groups displaying sub-species level genetic divergence similar to porpoises from the Black Sea. They are locally adapted to the upwelling environments off Iberia and NW Africa, with low population size and relatively isolated from neighboring populations (Fontaine, 2016; Fontaine et al., 2007; 2014; and see the assessment of the NAMMCO & IMR, 2019). Porpoises north of the Bay of Biscay belong to the *P. p. phocoena* subspecies. The very strong IBD detected at the mtDNA and, to a lesser extent, at microsatellite markers illustrate how geographically restricted dispersal is for this species. Local variation in IBD strength was reported here and previously (Fontaine et al., 2007) and reflect local variation in census size, dispersal behavior, and local habitat variation. Therefore, demes of the *P. p. phocoena* subspecies across the North Atlantic are far from a simple random mating unit. The strength of the IBD and its local variation suggest that local demes display variable demographic properties across the subspecies range, which is a function of local individual density and connectivity among neighboring demes. They are thus likely differentially impacted by fisheries bycatch. They should thus certainly be treated as distinct MUs for conservation and management assessments. The impact of bycatch in such a continuous stepping stone system under IBD is still poorly understood. It would require an in-depth treatment using simulations, modelling this system with realistic demographic parameters informed by direct (e.g., field survey and satellite tracking) or indirect estimations such as those provided by population genetics. This would allow assessing the resilience of harbor porpoises in the face of observed bycatch rates. Such simulations would be a valuable tool to assess the status of local demes across the North Atlantic, to design tailored conservation strategies, and identify areas of the species distribution range that are at high risk of local depletion that could disrupt gene flow and fragment populations.

All the individuals used in this study were sampled between 1990 and 2000. The genetic structure depicted here may thus just represent a temporal snapshot and may have already changed with the accelerated environmental changes we are experiencing. Ongoing environmental changes for instance have triggered profound reorganization of the North Atlantic ecosystems with impacts first upon primary production and in turn influenced top predators including harbor porpoises (Evans & Waggitt, 2020; Heide-Jørgensen et al., 2011; Lambert et al., 2014; MacLeod et al., 2005). Depletion of prey stocks induced by overfishing (Worm & Lotze, 2009) and the high rate of bycatch (NAMMCO & IMR, 2019; Orphanides & Palka, 2013) may have fostered habitat fragmentation and impacted the distribution of harbor porpoises, which could ultimately constitute a major threat for this small cetacean (Braulik et al., 2020; Evans, 2020; NAMMCO & IMR, 2019). It is thus paramount to reassess harbor porpoise genetic spatial structure with more recent samples to evaluate how it has evolved over the past 30 years. Finally, the pattern underlined in this study will provide key insights useful for devising management plans and model the future evolution of harbor porpoises with the forecasted climate changes. The AquaMap model predictions for the year 2050 under the most aggressive model (Figure 6, S16 to S18) show that abiotic environmental suitability for this particular species will not change dramatically compared to the present one. However, this model does not account for change in primary production and shifts in prey distributions. Porpoises are already impacted by climate change through the redistribution of their prey, as already observed (Evans & Waggitt, 2020; Hammond et al., 2002; Hammond et al., 2013; Heide-Jørgensen et al., 2011; Lambert et al., 2014; Mahfouz et al., 2017; NAMMCO & IMR, 2019).

The joint IMR-NAMMCO workshop on the status of harbor porpoises (2019) concluded that despite the absence of clear genetic discontinuities between porpoise demes across the *P. p. phocoena* distribution, pragmatic management units should be implemented using local connectivity assessment among areas and ecological and demographic information, while accounting for the possibility for porpoises to mix across certain units. This workshop (NAMMCO & IMR, 2019) also stressed the need for international collaborations to efficiently monitor porpoises over their distribution range and to continuously reassess these management units as the population structure highlighted here and therefore the suitable conservation measures might change in the future.

## Conclusion (294)

In the present study, we confirmed the undoubtedly deep separation existing between Pacific and Atlantic harbor porpoises that was previously identified as well as the acknowledged distinct ecotypes or subspecies of harbor porpoises previously reported in the Northeast Atlantic, Iberian and NW African upwelling waters, and in the Black Sea. However, we also discovered a new divergent mitochondrial lineage in one individual from West Greenland waters suggesting that a fourth ecotype may exist. It may be related to the new oceanic group of porpoises recently identified in that area which could have emerged during the LGM in an offshore glacial refugium (*e*.*g*. Azorean waters). These distinct ecotypes likely display specific adaptations, as suggested by their distinct behaviors, feeding ecology, habitat use, and genetic ancestries. Future genomic studies will certainly refine the evolutionary history of each ecotype and shed light on how natural selection may have contributed to their distinctiveness (Cammen et al., 2016). Besides additional genomic studies, knowledge about their habitat use, foraging preferences, demography, morphology, life history and behaviors is dearly required. Approaches such as niche habitat modelling, fatty acid and stable isotope analyses and satellite tracking should be implemented on each ecotype or subspecies to better grasp the nature of their differentiation and evolutionary trajectories. Our study suggests that harbor porpoise subpopulations from Northwest to Northeast Atlantic waters north of Biscay form a highly interconnected system without any major genetic discontinuities. However, it is important to realize that this large-scale continuous system is not a panmictic unit and in terms of conservation should be managed accordingly. The results of the present study complemented with further research on harbor porpoise life history, demography and ecology are critical to formulate management plans and improve the monitoring of the North Atlantic harbor porpoise in the context of the current climate crisis.

## Acknowledgements

We thank all the people involved in the sample collection, including KA Tolley, M Ferreira, T Jauniaux, J Haelters, AA Llavona, B and AA Ozturk, V Ridoux, F Caurant, W Dabin, E Rogan, M Sequeira, U Siebert, GA Vikingsson, A Borrell, AA Aguilar, D Palka, F Wenzel, M-P Heide-Jorgensen, AJ Read, G Stenson, V. Lesage, S. Barco and the national stranding networks: Réseau National d’Échouage and PELAGIS in France; Royal Belgian Institute of Natural Sciences-MUMM, and Dept. Pathology of the U. Liège, Belgium; CEMMA-Coordinadora para o Estudio dos Mamiferos Mariños, Spain; the Portuguese Marine Animal Tissue Bank-MTAB, Portugal; the Dutch Stranding Network, SOS Dolfijn, and Dept. Pathology of the U. Utrecht in the Netherlands; the Dept. pathology of the U. Kiel, Germany. We also thank Jeanine L. Olsen, Peter Evans, and the participants of the IMR-NAMMCO workshop in Tromsø (Norway, Dec. 2018) for their significant inputs during the development of this study. We are also grateful to the Centre for Information Technology of the University of Groningen, and in particular Bob Dröge and Fokke Dijkstra, for their continuous support and for providing access to the *Peregrine* high-performance computing cluster. This study benefitted was funded by the University of Groningen (The Netherlands). YBC was supported by a PhD fellowship from the University of Groningen.

## Data Availability

The microsatellite and mtDNA data supporting this study are available in the IRD Porpoises genetics and genomics Dataverse repository (https://dataverse.ird.fr/dataverse/porpoise_genomics) at doi: to be announced. Mitochondrial haplotypes were also deposited on Genbank under the accession numbers listed in Table S3.

## Author Contributions

MCF designed and supervised the study; PR coordinated the sampling in NWA and made them available for the present study; JT and RL performed the laboratory work and collected the data with help from MCF; YBC and RL analyzed the data; KK and CG generated the habitat models with help from YBC and MCF; YBC, RL and MCF interpreted the results and wrote the manuscript with input and final approval from all the co-authors.

