## Supplementary Information for "No leading-edge effect in North Atlantic harbor porpoises: Evolutionary and conservation implications"

### Tables

**Table S1.** Sampling stratified by geographical regions.

| Label | #individuals (mtDNA) | #individuals (microsatellites) | Sampling method | Origin |
|---|---|---|---|---|
| US | 18 | 71 | By-caught / Stranded | (Rosel et al., 1999) |
| CA | 5 | 60 | By-caught / Stranded | (Rosel et al., 1999) |
| WGLD | 28 | 29 | By-caught | (Rosel et al., 1999) |
| IC | 6 | 194 | By-caught | (Fontaine et al., 2014) |
| NN | 7 | 81 | Stranded | (Fontaine et al., 2014) |
| NS | 13 | 307 | Stranded | (Fontaine et al., 2014) |
| NBB | 15 | 58 | Stranded | (Fontaine et al., 2014) |
| IB | 18 | 33 | Stranded | (Fontaine et al., 2014) |
| MA | 14 | 14 | Stranded | (Fontaine et al., 2014) |
| BS | 12 | 78 | Stranded | (Fontaine et al., 2014) |

BS=Black Sea. MA=Mauritania. IB=Iberia. NBB=North Bay of Biscay. NS=North Sea. NN= North Norway. IC=Iceland. WGLD=West Greenland. CA= Canada. US= United states.

**Table S2.** Genetic diversity at the 10 microsatellites loci within the 30 geographical sub-groups.

| Populations | Subgroups | *N* | *Ho* | *He* | $F_{IS}$ | *Ar*‡ | *pAr*‡ |
|---|---|---|---|---|---|---|---|
| BS | BSBULG | 10.0 | 0.46 | 0.43 | -0.0798 | 1.45±0.08 | 0.03±0.02 |
| | BSGEO | 8.0 | 0.56 | 0.47 | -0.2000 | - | - |
| | BSTK | 14.6 | 0.44 | 0.47 | 0.0537 | 1.49±0.07 | 0.04±0.01 |
| | BSTKM | 2.9 | 0.52 | 0.37 | -0.3933 | - | - |
| | BSU | 42.0 | 0.51 | 0.49 | -0.0400 | 1.49±0.07 | 0.04±0.01 |
| MA | MA | 12.9 | 0.61 | 0.61 | -0.0128 | 1.63±0.05 | 0.06±0.03 |
| IB | PT | 12.8 | 0.64 | 0.59 | -0.0932 | 1.61±0.06 | 0.08±0.03 |
| | SP | 18.9 | 0.51 | 0.52 | 0.0283 | 1.54±0.07 | 0.08±0.04 |
| NBB | BB | 25.7 | 0.69 | 0.72 | 0.0332 | 1.73±0.05 | 0.08±0.02 |
| | IRCS | 27.9 | 0.69 | 0.73 | 0.0457 | 1.74±0.07 | 0.10±0.03 |
| NS | IRIS | 8.9 | 0.60 | 0.68 | 0.1192 | - | - |
| | IRAT | 7.0 | 0.74 | 0.71 | -0.0475 | - | - |
| | SC | 18.8 | 0.74 | 0.79 | 0.0571 | 1.81±0.05 | 0.12±0.02 |
| | FRC | 49.7 | 0.75 | 0.75 | -0.0044 | 1.76±0.07 | 0.08±0.02 |
| | BL | 54.5 | 0.76 | 0.76 | 0.0047 | 1.76±0.07 | 0.10±0.03 |
| | H | 30.9 | 0.77 | 0.78 | 0.0092 | 1.79±0.06 | 0.13±0.02 |
| | G | 45.5 | 0.72 | 0.77 | 0.0630 | 1.78±0.06 | 0.14±0.02 |
| | DK | 4.8 | 0.75 | 0.64 | -0.1630 | - | - |
| | IFR | 67.1 | 0.72 | 0.76 | 0.0526 | 1.76±0.08 | 0.13±0.04 |
| | N1 | 70.0 | 0.72 | 0.76 | 0.0499 | 1.77±0.08 | 0.12±0.03 |
| NN | N2 | 10.0 | 0.74 | 0.72 | -0.0292 | 1.77±0.07 | 0.10±0.02 |
| IC | ICN | 57.9 | 0.72 | 0.77 | 0.0632 | 1.78±0.06 | 0.11±0.02 |
| | ICSE | 14.0 | 0.78 | 0.74 | -0.0506 | 1.77±0.07 | 0.12±0.02 |
| | ICSW | 71.9 | 0.71 | 0.76 | 0.0676 | 1.77±0.08 | 0.10±0.02 |
| | ICW | 50.0 | 0.75 | 0.77 | 0.0209 | 1.76±0.08 | 0.09±0.02 |
| WGLD | WGLD | 29.0 | 0.76 | 0.75 | -0.0176 | 1.76±0.09 | 0.09±0.03 |
| CA | NF | 28.6 | 0.77 | 0.75 | -0.0176 | 1.76±0.07 | 0.11±0.03 |
| | SL | 28.0 | 0.77 | 0.76 | -0.0195 | 1.77±0.08 | 0.12±0.02 |
| US | GM | 13.8 | 0.52 | 0.63 | 0.1750* | 1.66±0.11 | 0.08±0.02 |
| | US | 49.3 | 0.74 | 0.75 | 0.0250 | 1.76±0.08 | 0.14±0.03 |

*N*, microsatellite averaged sample size; *He* and *Ho,* expected and observed heterozygosity; $F_{IS}$, fixation index; *Ar*, allelic richness; *PAr*, private allelic richness. Assuming a samples size of 14 haploid chromosomes (‡). *$p$-value ≤ 0.05. The populations with less than 10 individuals were discard from some analyses.

BSBULG= Black Sea Bulgaria. BSGEO= Black Sea Georgia. BSTKM=Black Sea Turkey Marmara Sea. BSTK=Black Sea Turkey. BSU=Black Sea Ukraine. MA=Mauritania. PT= Portugal. SP= Spain. BB=Bay of Biscay. IRCS= Celtic Sea. FRC=France Channel. IRIS=. Irish Sea. IRAT=Irish Atlantic. SC=Scotland. BL=Belgium. H=Holland. G=Germany. DK=Denmark. IFR=Faroe Island. N1=Norway South. N2=Norway North. ICN=Iceland North. ICSE=Iceland South East. ICSW=Iceland South West. ICW=Iceland West. WGLD=West Greenland. NF= Newfoundland. SL=Saint Lawrence. GM=Gulf of Maine. US=United States.

**Table S3.** Lists of mitochondrial haplotypes carried by each individual sorted per population.

| ID | Geo. Regions | Haplotype | Genbank Accession ID |
|---|---|---|---|
| B11 | BS | Hap 19 | |
| TK3 | BS | Hap 21 | |
| TK12 | BS | Hap 22 | |
| U30 | BS | Hap 23 | |
| U31 | BS | Hap 24 | |
| U75 | BS | Hap 24 | |
| U93 | BS | Hap 24 | |
| U64 | BS | Hap 25 | |
| U69 | BS | Hap 26 | |
| TK10 | BS | Hap 29 | |
| TK11 | BS | Hap 45 | |
| U49 | BS | Hap 46 | |
| 156 | MA | Hap 14 | |
| 88 | MA | Hap 14 | |
| 89 | MA | Hap 14 | |
| 153 | MA | Hap 15 | |
| 99 | MA | Hap 15 | |
| 106 | MA | Hap 16 | |
| 132 | MA | Hap 16 | |
| 100 | MA | Hap 17 | |
| 141 | MA | Hap 17 | |
| 109 | MA | Hap 18 | |
| 125 | MA | Hap 18 | |
| 189 | MA | Hap 18 | |
| 97 | MA | Hap 27 | |
| 140 | MA | Hap 28 | |
| PP79-2003 | IB | Hap 20 | |
| PP04-2002 | IB | Hap 30 | |
| PPH021 | IB | Hap 30 | |
| PPH027 | IB | Hap 30 | |
| PPH028 | IB | Hap 30 | |
| PPH029 | IB | Hap 30 | |
| PPH030 | IB | Hap 30 | |
| PPH032 | IB | Hap 30 | |
| PP29-2002 | IB | Hap 33 | |
| PP30-2002 | IB | Hap 34 | |
| PP63-2002 | IB | Hap 35 | |
| PP68-2002 | IB | Hap 36 | |
| PP79-2000 | IB | Hap 37 | |
| PP118-2004 | IB | Hap 38 | |
| PPH009 | IB | Hap 39 | |
| PPH017 | IB | Hap 40 | |
| PPH035 | IB | Hap 41 | |
| BS51 | IB | Hap 42 | |
| FR9904033 | NBB | Hap 10 | |
| FR10003091 | NBB | Hap 13 | |
| FR9712124 | NBB | Hap 30 | |
| FRPP4 | NBB | Hap 30 | |
| FR10304042 | NBB | Hap 31 | |
| FR10402044 | NBB | Hap 32 | |
| FR10003094 | NBB | Hap 43 | |
| FR9903024 | NBB | Hap 43 | |
| FR10007111 | NBB | Hap 44 | |
| FR10001003 | NBB | Hap 48 | |
| FR10002054 | NBB | Hap 48 | |
| FR10205213 | NBB | Hap 54 | |

| | | |
|---|---|---|
| FR10403052 | NBB | Hap 66 |
| FR10405060 | NBB | Hap 66 |
| FR9904040 | NBB | Hap 66 |
| IFR1 | NS | Hap 50 |
| IFR3 | NS | Hap 50 |
| IFR4 | NS | Hap 50 |
| IFR8 | NS | Hap 50 |
| IFR5 | NS | Hap 6 |
| IFR6 | NS | Hap 6 |
| 99-22 | NS | Hap 66 |
| 2000-32 | NS | Hap 68 |
| 2000-03 | NS | Hap 7 |
| 2000-50 | NS | Hap 70 |
| 2000-04 | NS | Hap 79 |
| 2000-26 | NS | Hap 8 |
| IFR10 | NS | Hap 9 |
| VF-05-99 | NN | Hap 5 |
| 2000-10 | NN | Hap 55 |
| 2000-24 | NN | Hap 65 |
| 99-24 | NN | Hap 67 |
| 2000-38 | NN | Hap 69 |
| 2000-44 | NN | Hap 71 |
| 2000-41 | NN | Hap 86 |
| H162 | IC | Hap 10 |
| SV226 | IC | Hap 11 |
| H139 | IC | Hap 49 |
| AK31 | IC | Hap 56 |
| SV253-92 | IC | Hap 62 |
| H211 | IC | Hap 78 |
| 803 | WGLD | Hap 2 |
| 772 | WGLD | Hap 3 |
| 816 | WGLD | Hap 3 |
| 801 | WGLD | Hap 4 |
| 778 | WGLD | Hap 47 |
| 771 | WGLD | Hap 5 |
| 791 | WGLD | Hap 52 |
| 769 | WGLD | Hap 57 |
| 761 | WGLD | Hap 58 |
| 753 | WGLD | Hap 59 |
| 777 | WGLD | Hap 61 |
| 774 | WGLD | Hap 63 |
| 817 | WGLD | Hap 64 |
| 796 | WGLD | Hap 73 |
| 797 | WGLD | Hap 76 |
| 759 | WGLD | Hap 77 |
| 764 | WGLD | Hap 77 |
| 811 | WGLD | Hap 80 |
| 828 | WGLD | Hap 82 |
| 783 | WGLD | Hap 84 |
| 757 | WGLD | Hap 87 |
| 819 | WGLD | Hap 89 |
| 751 | WGLD | Hap 90 |
| 823 | WGLD | Hap 93 |
| 827 | WGLD | Hap 93 |
| 807 | WGLD | Hap 94 |
| 815 | WGLD | Hap 97 |
| 813 | WGLD | Hap 98 |
| Pp210801B | CA | Hap 12 |
| Pp130701 | CA | Hap 84 |
| F20020412 | CA | Hap 88 |
| Pp030702 | CA | Hap 92 |

| | | |
|---|---|---|
| Pp190701B | CA | Hap 99 |
| VAQS20131038 | US | Hap 1 |
| VAQS20131039 | US | Hap 51 |
| VAQS20131011 | US | Hap 53 |
| VGT258 | US | Hap 60 |
| VAQS20131033 | US | Hap 72 |
| VAQS20101011 | US | Hap 74 |
| VAQS20141003 | US | Hap 75 |
| VAQS20131027 | US | Hap 81 |
| VAQS20131030 | US | Hap 83 |
| VAQS20101018 | US | Hap 84 |
| VAQS20121015 | US | Hap 84 |
| VAQS20131005 | US | Hap 84 |
| VAQS20131028 | US | Hap 84 |
| VGT264 | US | Hap 84 |
| VAQS20131046 | US | Hap 85 |
| KLC098 | US | Hap 91 |
| CALO1203 | US | Hap 95 |
| VAQS20141005 | US | Hap 96 |
| 161 | NP | Hap 101 |
| 706 | NP | Hap 102 |
| 3767 | NP | Hap 102 |
| 1082 | NP | Hap 102 |
| 1080 | NP | Hap 103 |
| 4764 | NP | Hap 104 |
| 26606 | NP | Hap 105 |
| 5411 | NP | Hap 106 |
| 3768 | NP | Hap 107 |
| 1084 | NP | Hap 108 |
| 8514 | NP | Hap 109 |
| 704 | NP | Hap 110 |
| 3766 | NP | Hap 111 |
| 707 | NP | Hap 112 |

**Table S4.** Genetic diversity at the mtDNA markers.

| **<u>mtDNA</u>** | **All individuals** | ***P. p. vomerina*** | ***P. p. phocoena*** | | | | | | | ***Hybrids*♠** | ***IBMA*** *(P. p. meridionalis)* | | ***P. p. relicta*** |
|---|---|---|---|---|---|---|---|---|---|---|---|---|---|
| | All | NP | All North Atlantic (NAT) | US | CA | WGLD† | IC | NN | NS | NBB | IB | MA | BS |
| $N_{mtDNA}$ | 148 | 12 | 76 | 18 | 5 | 27 | 6 | 7 | 13 | 15 | 18 | 14 | 12 |
| *S* | 359 | 35 | 196 | 73 | 28 | 111 | 40 | 37 | 57 | 71 | 17 | 12 | 26 |
| *Singl.* | 118 | 15 | 100 | 50 | 24 | 67 | 35 | 29 | 24 | 21 | 13 | 1 | 24 |
| Parsim. | 241 | 20 | 96 | 23 | 4 | 44 | 5 | 8 | 33 | 50 | 4 | 11 | 2 |
| *# hap* | 110 | 11 | 62 | 14 | 5 | 24 | 6 | 7 | 9 | 10 | 12 | 7 | 10 |
| *# priv. hap* | | 11 | 57 | 13 | 4 | 22 | 5 | 6 | 8 | 8 | 11 | 7 | 10 |
| *Hd* | 0.99 | 0.99 | 0.99 | 0.94 | 1.00 | 0.99 | 1.00 | 1.00 | 0.91 | 0.94 | 0.86 | 0.90 | 0.96 |
| $\pi$ (%) | 0.80 | 0.21 | 0.36 | 0.31 | 0.27 | 0.37 | 0.33 | 0.29 | 0.38 | 0.52 | 0.05 | 0.07 | 0.10 |
| $\pi$ (%)✯ | 0.76 | 0.21 | 0.36 | 0.31 | 0.27 | 0.37 | 0.33 | 0.29 | 0.38 | 0.52 | 0.05 | 0.07 | 0.10 |
| $\theta_W$ (%) | 1.15 | 0.27 | 0.90 | 0.48 | 0.30 | 0.65 | 0.39 | 0.34 | 0.41 | 0.49 | 0.11 | 0.09 | 0.19 |
| $\theta_W$ (%)✯ | 0.84 | 0.22 | 0.39 | 0.34 | 0.30 | 0.40 | 0.37 | 0.31 | 0.39 | 0.51 | 0.06 | 0.07 | 0.12 |
| *D* | – | – | – | -1.38 | -0.79 | -1.69 | -1.08 | -0.85 | -0.30 | 0.25 | -2.00* | -0.66 | -2.07** |
| *D*±SE✯ | – | – | – | -0.39±0.55 | -0.79±0.00 | -0.60±0.42 | -0.85±0.06 | -0.49±0.29 | -0.07±0.64 | 0.24±0.75 | -0.96±0.26 | -0.48±0.77 | -1.11±0.18 |
| *D** | – | – | – | -2.11* | -0.79 | -2.33* | -1.15 | -1.07 | -0.34 | 0.17 | -2.55* | 1.1 | -2.46** |
| *D**±SE✯ | – | – | – | -0.43±0.53 | -0.79±0.00 | -0.60±0.41 | -0.85±0.06 | -0.50±0.30 | -0.07±0.65 | 0.20±0.79 | -0.97±0.24 | -0.47±0.78 | -1.12±0.17 |

$N_{mtDNA}$, mtDNA sample sizes; *S*, segregating sites; *Singl.*, singleton; Parsim., sites informative in parsimony; *# hap*, number of haplotypes; *# priv. hap*, number of private haplotypes; *Hd*, haplotype diversity; $\pi$, nucleotide diversity; $\theta_W$, Watterson's theta; *D*, Tajima's *D*; *D**, Fu and Li's D*; Assuming a samples size of 5 individuals (✯). NS: not significant (*p*-value > 0.05); Hybrid lineage showing an admix genetic ancestry from *P. p. meridionalis* and *P. p. phocoena* (♠) **p*-value ≤ 0.05; ***p*-value ≤ 0.01; ****p*-value ≤ 0.001. BS=Black Sea. MA=Mauritania. IB=Iberia. NBB=North Bay of Biscay. NS=North Sea. NN=North Norway. IC=Iceland. WGLD=West Greenland. CA=Canada. US=United states.

**Table S5.** Genetic diversity at the 10 microsatellite loci

| | $N_{Msat}$ | *He* | *Ho* | $F_{IS}$ | *Ar* ± SE‡ | *PAr* ± SE‡ | $M_{GW}$ ± SE |
|---|---|---|---|---|---|---|---|
| ***P. p. phocoena*** | | | | | | | |
| **All (NAT)** | 705.8 | 0.77 | 0.73 | $0.046^{*}$ | 7.21 ± 0.89 | – | 0.46 ± 0.07 |
| **US** | 63.1 | 0.75 | 0.7 | $0.066^{NS}$ | 7.26 ± 0.97 | 0.51 ± 0.15 | 0.46 ± 0.07 |
| **CA** | 56.6 | 0.76 | 0.77 | $-0.010^{NS}$ | 7.19 ± 0.94 | 0.30 ± 0.05 | 0.49 ± 0.09 |
| **WGLD**† | 29 | 0.75 | 0.76 | $-0.018^{NS}$ | 7.19 ± 0.95 | 0.23 ± 0.06 | 0.52 ± 0.05 |
| **IC** | 193.8 | 0.77 | 0.73 | $0.056^{*}$ | 7.14 ± 0.88 | 0.31 ± 0.04 | 0.47 ± 0.04 |
| **NN** | 70 | 0.76 | 0.72 | $0.050^{*}$ | 7.14 ± 0.93 | 0.27 ± 0.07 | 0.49 ± 0.09 |
| **NS** | 293.3 | 0.77 | 0.74 | $0.040^{*}$ | 7.20 ± 0.87 | 0.39 ± 0.05 | 0.44 ± 0.07 |
| ***Hybrids*♠** | | | | | | | |
| **NBB** | 57.5 | 0.73 | 0.69 | $0.047^{NS}$ | 6.47 ± 0.87 | 0.27 ± 0.07 | 0.44 ± 0.06 |
| ***IBMA*** *(P. p. meridionalis)* | | | | | | | |
| **IB** | 31.7 | 0.56 | 0.56 | $-0.008^{NS}$ | 4.42 ± 0.72 | 0.26 ± 0.09 | 0.30 ± 0.03 |
| **MA** | 12.9 | 0.61 | 0.61 | $-0.013^{NS}$ | 4.53 ± 0.75 | 0.20 ± 0.13 | 0.35 ± 0.00 |
| ***P. p. relicta*** | | | | | | | |
| **BS** | 77.5 | 0.49 | 0.5 | $-0.015^{NS}$ | 3.49 ± 0.43 | 0.24 ± 0.06 | 0.30 ± 0.05 |

$N_{Msat}$, microsatellite averaged sample size; *He* and *Ho*, expected and observed heterozygosity; $F_{IS}$, fixation index; *Ar*, allelic richness; *PAr*, private allelic richness; $M_{GW}$, M-ratio. † Excluding Hap 47 for mtDNA. ‡ Assuming a sample size of 18 haploid chromosomes. ♠ Hybrid group as previously identified in Fontaine et al. (2014 and 2017) showing admixed ancestry in the STRUCTURE analyses between *P. p. meridionalis* and *P. p. phocoena.* NS: not significant (*p*-value > 0.05); **p*-value ≤ 0.05; ***p*-value ≤ 0.01; ****p*-value ≤ 0.001. BS=Black Sea. MA=Mauritania. IB=Iberia. NBB=North Bay of Biscay. NS=North Sea. NN=North Norway. IC=Iceland. WGLD=West Greenland. CA=Canada. US=United states

**Table S6.** Estimated effective population size (*Ne*) within each geographic region estimated using *NeEstimator* (Do et al., 2014). *Ne* values are provided as median and 95% confidence interval (CI) of diploid individuals (See table S1for the acronyms).

| Population | Median Ne | 95% CI |
|---|---|---|
| **BS** | 504 | [133-1,120] |
| **MA** | 12 | [6-30.9] |
| **IB** | 56 | [30-173] |
| **NBB** | 217 | [121-788] |
| **NS** | 1,777 | [662-1,077] |
| **NN** | 2,882 | [358-∞] |
| **IC** | 819 | [489-2,190] |
| **WGLD** | 925 | [119-∞] |
| **CA** | 1,482 | [391-∞] |
| **US** | 2,624 | [244-∞] |

**Table S7.** Per generation migration rate estimates. Statistically significantly asymmetric values ($p$-value < 0.05) are indicated with an asterisk. (See table S1for the acronyms).

| From \ To | BS | MA | IB | NBB | NS | NN | IC | WGLD | CA | US |
|---|---|---|---|---|---|---|---|---|---|---|
| **BS** | - | 0.001 | 0.001 | 0.001 | 0.001 | 0.000 | 0.000 | 0.001 | 0.001 | 0.000 |
| **MA** | 0.001 | - | 0.021 | 0.057 | 0.008 | 0.003 | 0.002 | 0.003 | 0.002 | 0.002 |
| **IB** | 0.000 | 0.009 | - | 0.064 | 0.024* | 0.006 | 0.000 | 0.004 | 0.008 | 0.001 |
| **NBB** | 0.001 | 0.021 | 0.026 | - | 0.152 | 0.174 | 0.142 | 0.083 | 0.112 | 0.098 |
| **NS** | 0.000 | 0.000 | 0.000* | 0.160 | - | 0.715 | 1 | 0.251 | 0.492 | 0.454 |
| **NN** | 0.000 | 0.000 | 0.000 | 0.156 | 0.915 | - | 0.578 | 0.270 | 0.431 | 0.362 |
| **IC** | 0.000 | 0.000 | 0.000 | 0.143 | 0.976 | 0.190 | - | 0.246 | 0.608 | 0.345 |
| **WGLD** | 0.000 | 0.000 | 0.000 | 0.111 | 0.368 | 0.274 | 0.317 | - | 0.214 | 0.284 |
| **CA** | 0.000 | 0.000 | 0.000 | 0.134 | 0.618 | 0.458 | 0.624 | 0.177 | - | 0.309 |
| **US** | 0.000 | 0.000 | 0.000 | 0.107 | 0.654 | 0.422 | 0.490 | 0.239 | 0.327 | - |

**Table S8.** Effective number of migrants (*2.Ne.m*) per generation.

| From \ To | BS | MA | IB | NBB | NS | NN | IC | WGLD | CA | US |
|---|---|---|---|---|---|---|---|---|---|---|
| **BS** | - | 0 | 0 | 1 | 0 | 0 | 0 | 0 | 0 | 0 |
| **MA** | 0 | - | 0 | 1 | 0 | 0 | 0 | 0 | 0 | 0 |
| **IB** | 0 | 0 | - | 7 | 0 | 0 | 0 | 0 | 0 | 0 |
| **NBB** | 0 | 8 | 11 | - | 66 | 75 | 61 | 35 | 48 | 42 |
| **NS** | 0 | 0 | 0 | 567 | - | 2540 | 3554 | 893 | 1748 | 1612 |
| **NN** | 0 | 0 | 0 | 898 | 5272 | - | 3329 | 1553 | 2486 | 2084 |
| **IC** | 0 | 0 | 0 | 234 | 1599 | 310 | - | 403 | 995 | 565 |
| **WGLD** | 0 | 0 | 0 | 204 | 680 | 506 | 586 | - | 396 | 525 |
| **CA** | 0 | 0 | 0 | 396 | 1831 | 1358 | 1848 | 524 | - | 914 |
| **US** | 0 | 0 | 0 | 562 | 3433 | 2213 | 2572 | 1256 | 1718 | - |

(See table S1for the acronyms).

**Table S9**. Environmental envelope settings used for AquaMaps species distribution modelling.

| | Min | Pref. Min (10th percentile) | Pref. Max (90th percentile) | Max |
|---|---|---|---|---|
| Depth (m) | 0 | 8 | 282 | 3000 |
| Temperature (ºC) | -0.37 | 7.35 | 15.31 | 23 |
| Salinity (psu) | 3.61 | 19.62 | 35.27 | 40 |
| Sea Ice Concentration | -1 | 0 | 0 | 0.2 |

### Figures

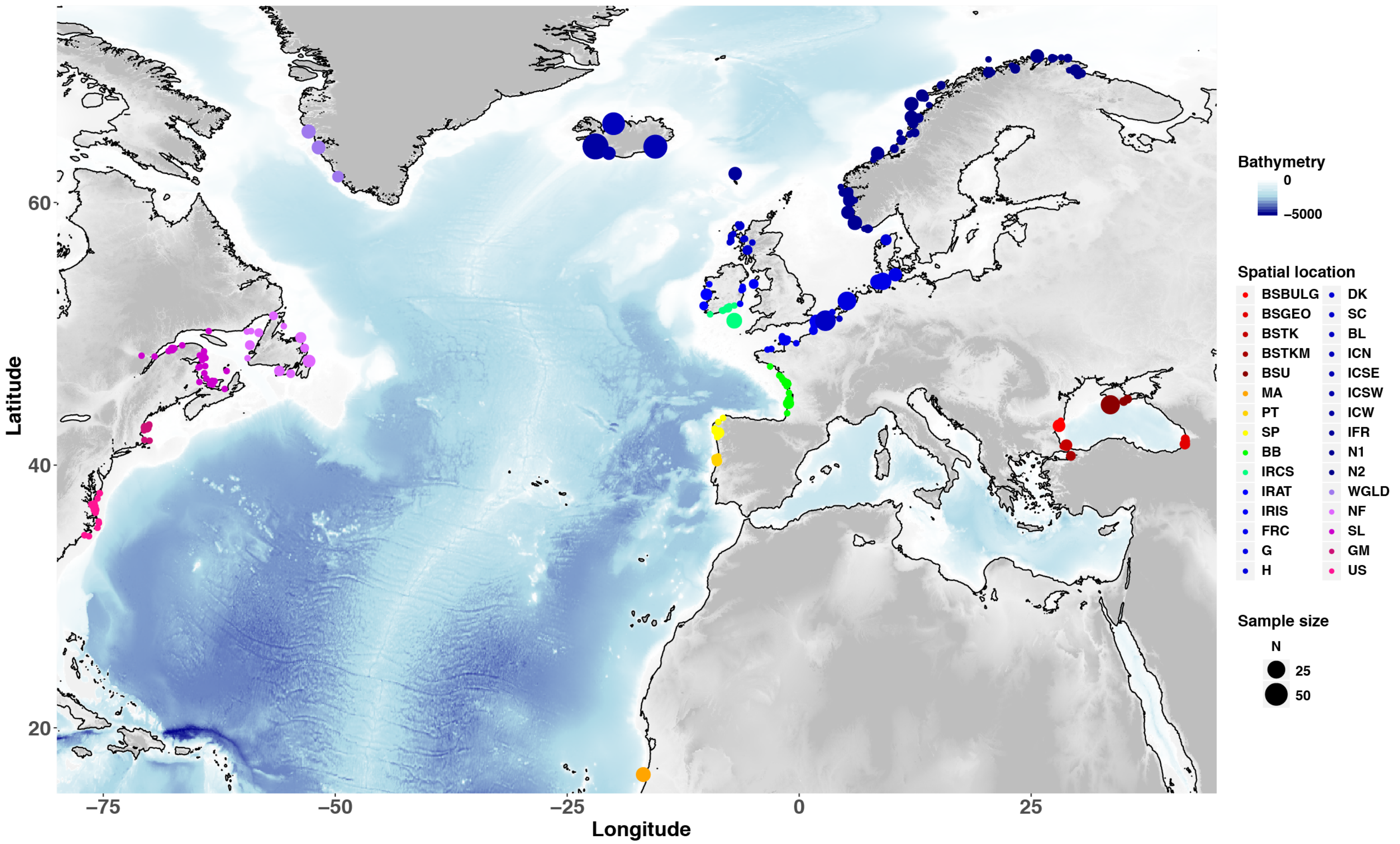

**Figure S1.** Map showing the sampling locations and geographic partitioning of the 30 geographical sub-groups considered for microsatellites data analyses at the finest scale. Sampling locations are based on approximate GPS coordinates or reported discovery location. (See Table S2 for the acronyms).

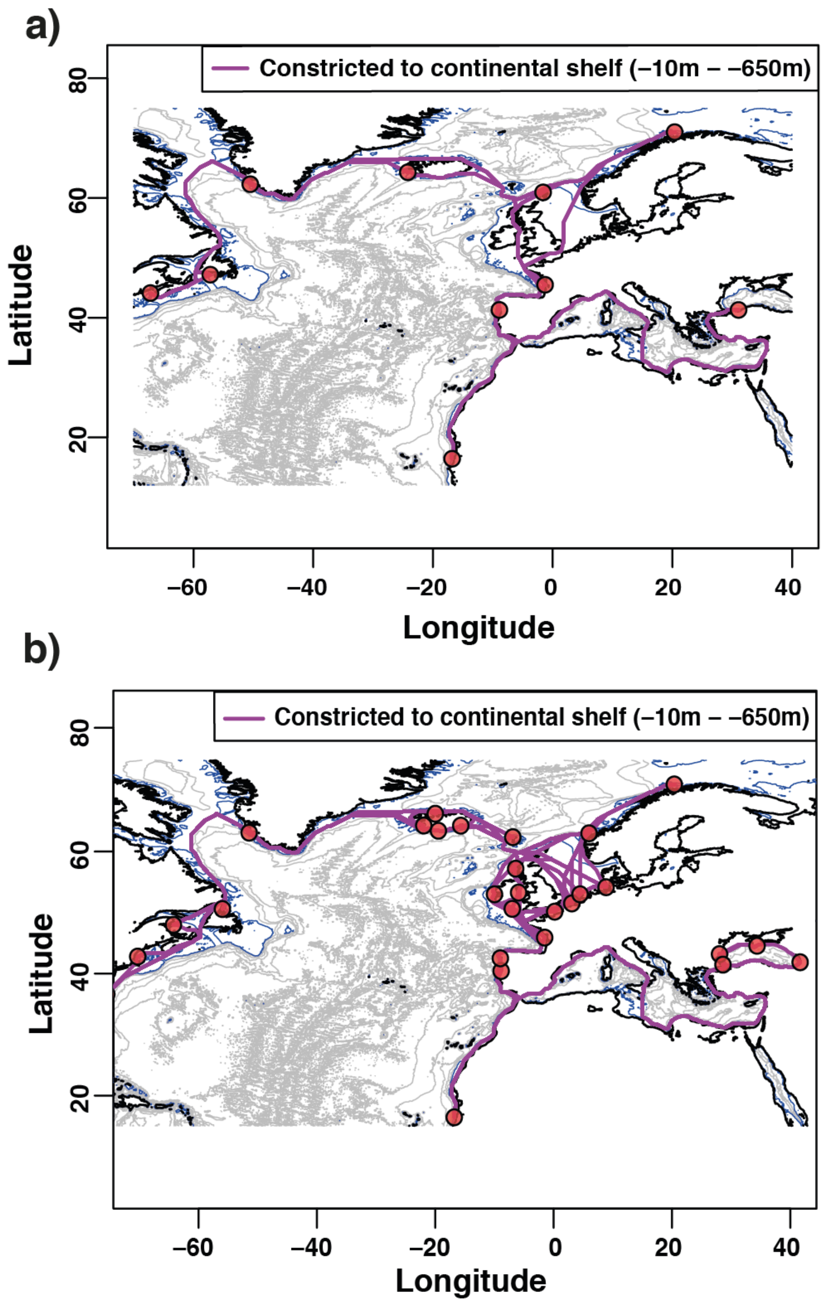

**Figure S2.** Map showing the marine distances restricting porpoise movements between -10m and -650m among (a) the 10 geographical regions and (b) the 30 subgroups.

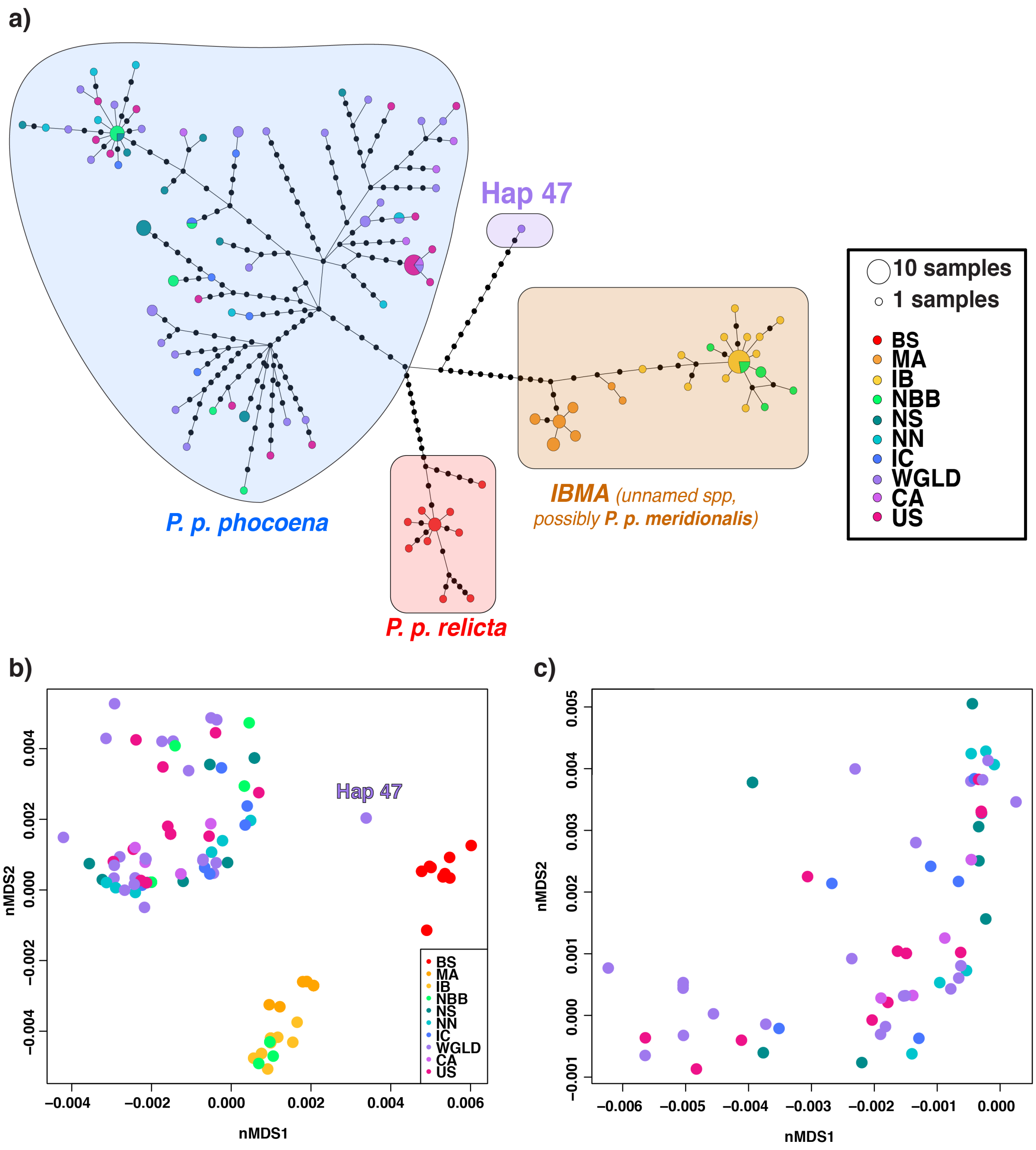

**Figure S3.** Phylogenetic relationships among mtDNA haplotypes represented using a median-joining (MJ) haplotype network (a) and a non-metric multidimensional scaling (nMDS) (b,c).

**(a)** Mitochondrial MJ haplotype network for samples in the Atlantic Ocean and Black Sea. Each circle represents a haplotype. The circle sizes are proportional to the haplotype frequencies observed in the total sampling, and each pie slice is proportional to the haplotype counts observed per geographic location. Black nodes represent mutational steps between haplotypes.

**(b and c)** NMDS describing mtDNA genetic relationships based on the Jukes-Cantor distance between pair of sequences along the first two axes for **(b)** the 10 geographical regions and **(c)** focusing only on the North Atlantic individuals (*P. p. phocoena*) excluding Hap 47. See Table S1 for the group acronyms.

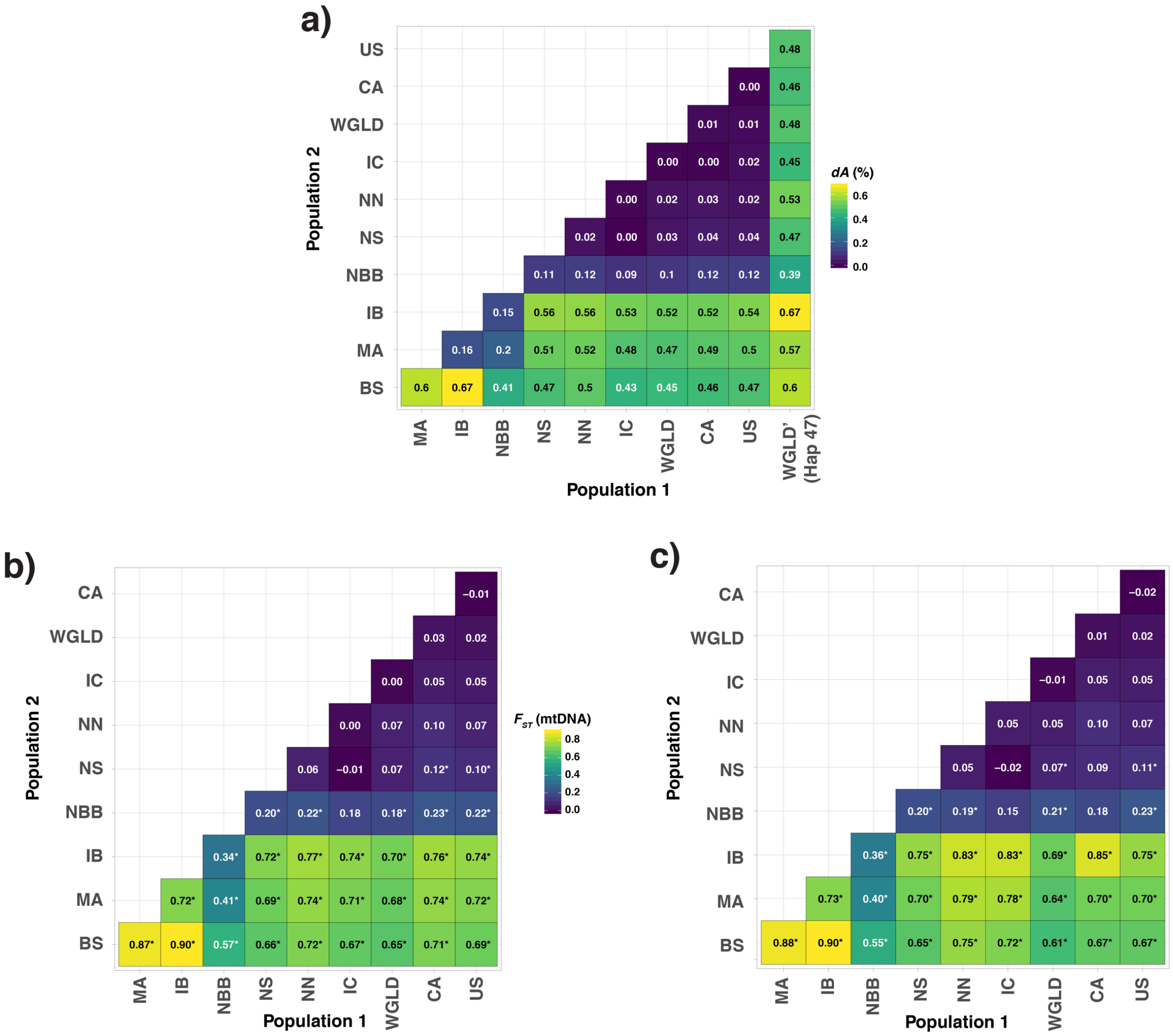

**Figure S4.** Pairwise mtDNA divergence **(a)** and differentiation **(b,c)** between geographical regions. Acronyms are presented in Table S1. WGLD' (Hap 47) represents the highly divergent haplotype found in WGLD and its level of divergence compared to the other mtDNA haplotypes (see Fig. 2, 3 and S3a). *p-value < 0.05. See Table S1 for the group acronyms.

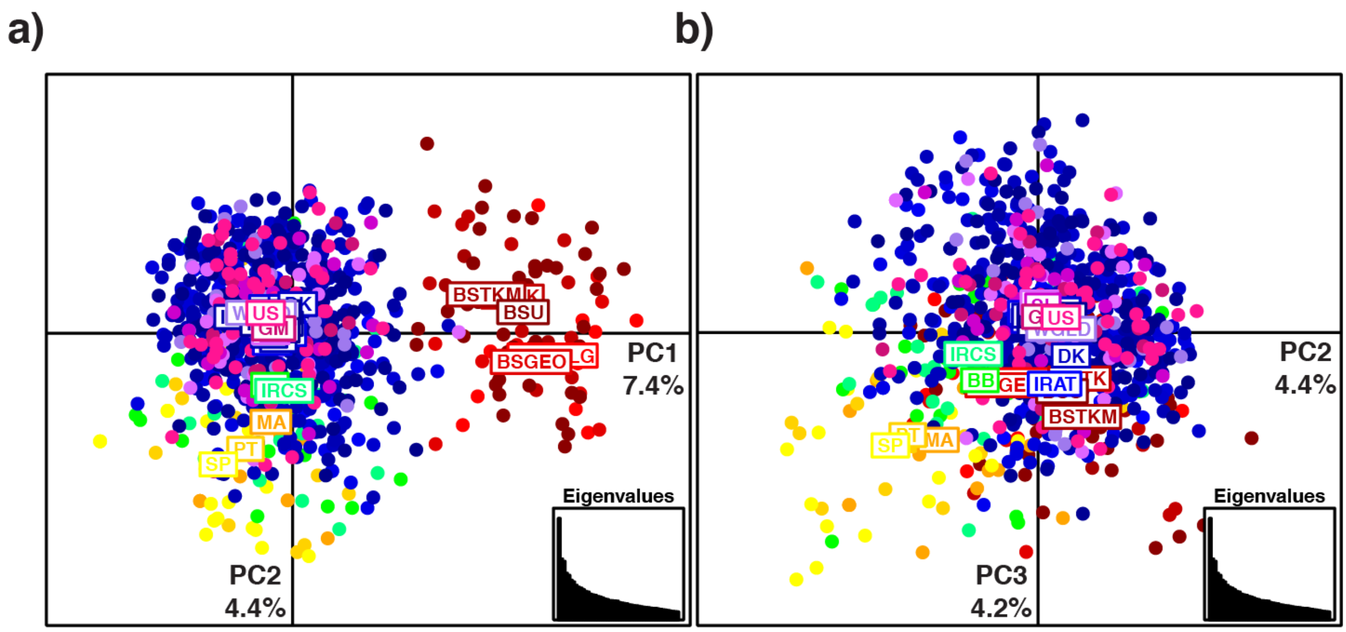

**Figure S5.** Principal components analysis (PCA) showing harbor porpoise genetic structure at the microsatellite markers. The scatter plots show the two first PCs **(a)** and the second and third PCs **(b)**. Points are color-coded according to the 30 geographical subgroups. See Table S2 for group acronyms.

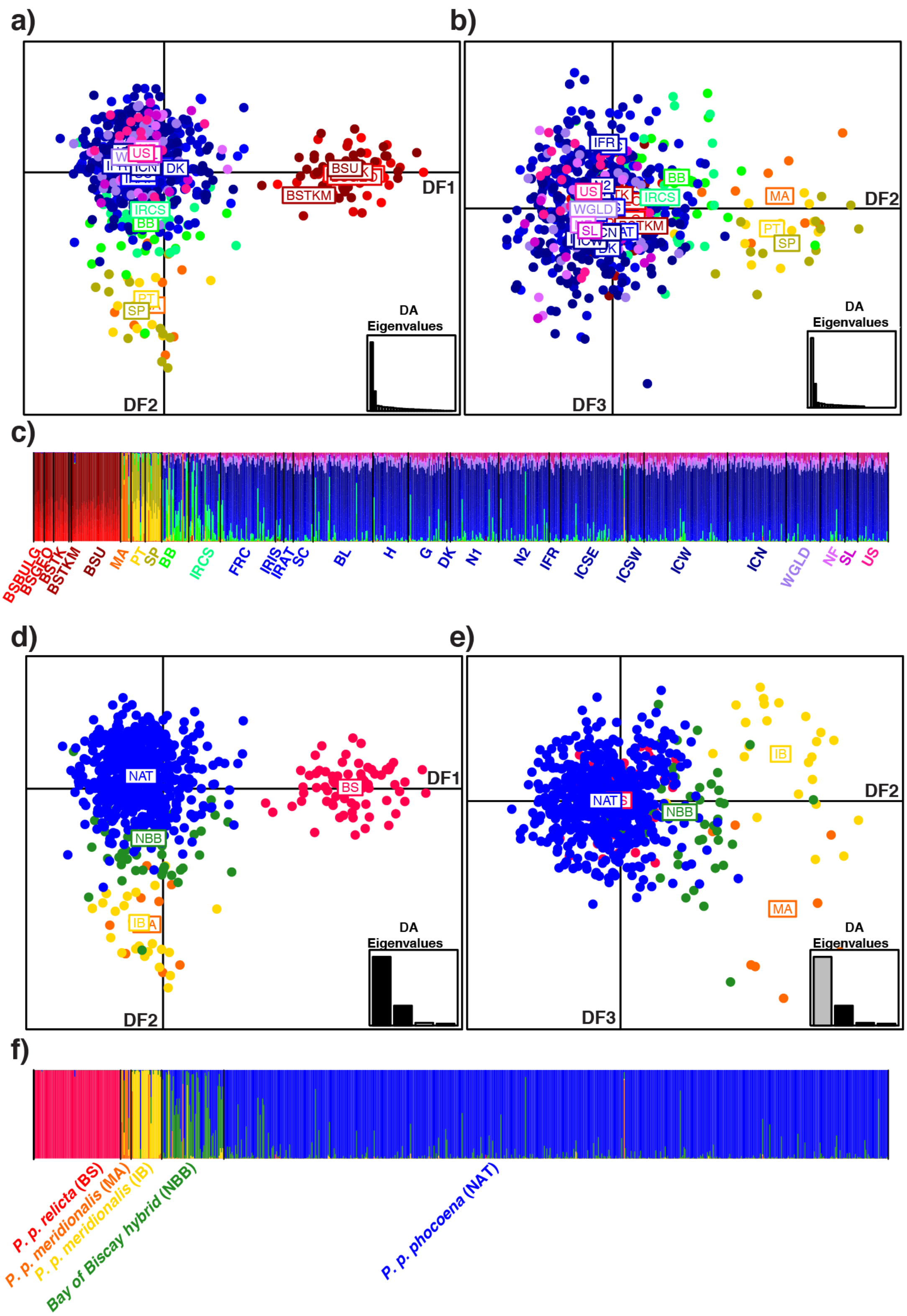

**Figure S6.** Discriminant analysis of principal components (DAPC) showing harbor porpoise genetic structure at the microsatellite markers based on two *a priori* grouping: (a–c) the 30 geographical subgroups (see Table S2 for accronyms) and (d–f) five groups including the four mtDNA lineages and sublineages, and the hybrids. For the two grouping, results are shown as scatter plots for the first two first discriminants functions (DFs; DF1 and DF2), and DF2 and DF3, as well as barplots showing the cluster membership probability of the DAPC. See Table S2 for acronyms.

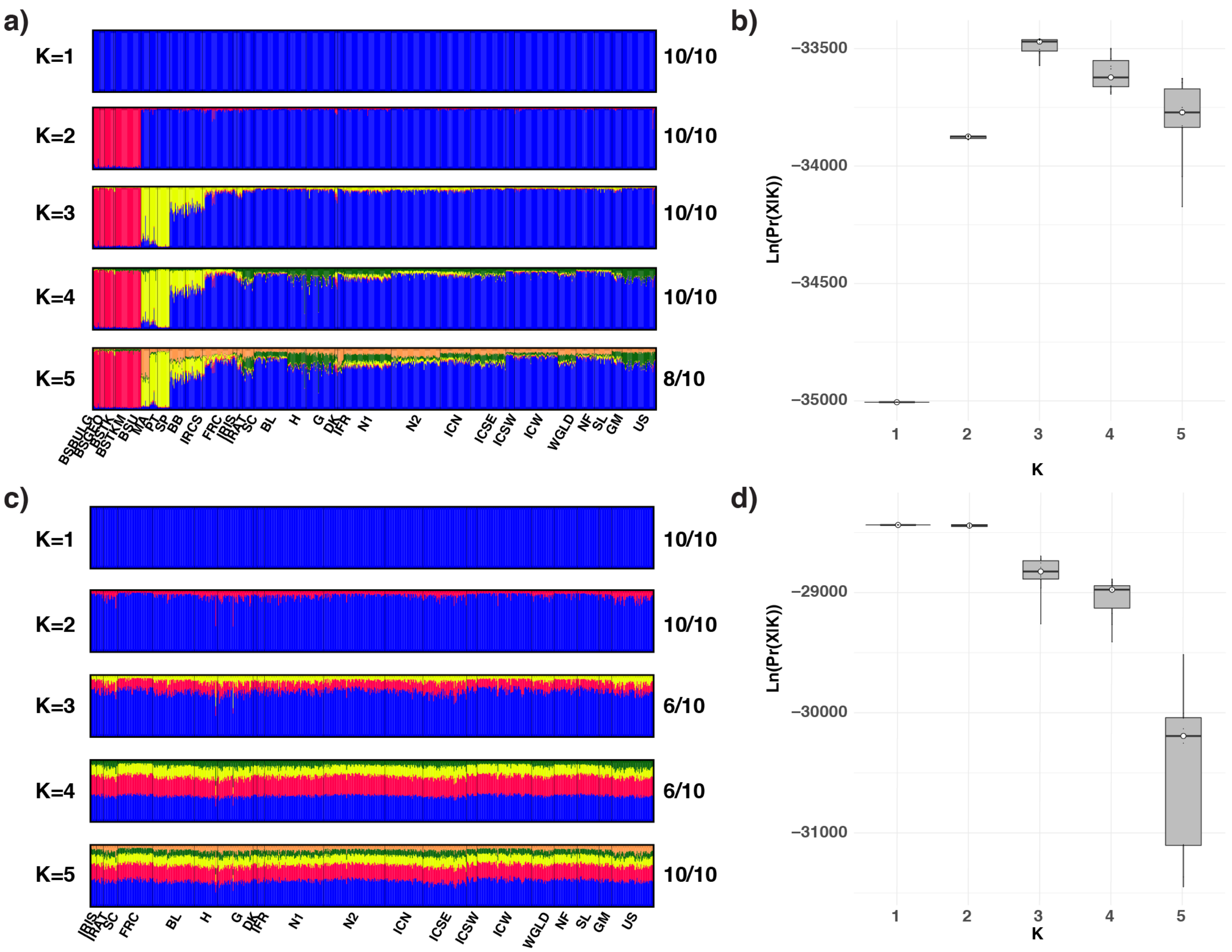

**Figure S7.** Population structure estimated using the Bayesian clustering approach of *STRUCTURE* including individuals with missing data. In the barplots (a and c) each individual is represented by a vertical line divided into *K* segments showing the genetic ancestry proportions to each of the *K* clusters for (a) the whole data set or (c) just for *P. p. phocoena* individuals in the North Atlantic. Numbers on the right side of the barplots show the number of times this result was found out of 10 independent replicated analyses. The estimated probability of the data (X) given *K* clusters tested is shown for the whole data set (b) and just for the North Atlantic individuals (d). Acronyms are presented in Table S2.

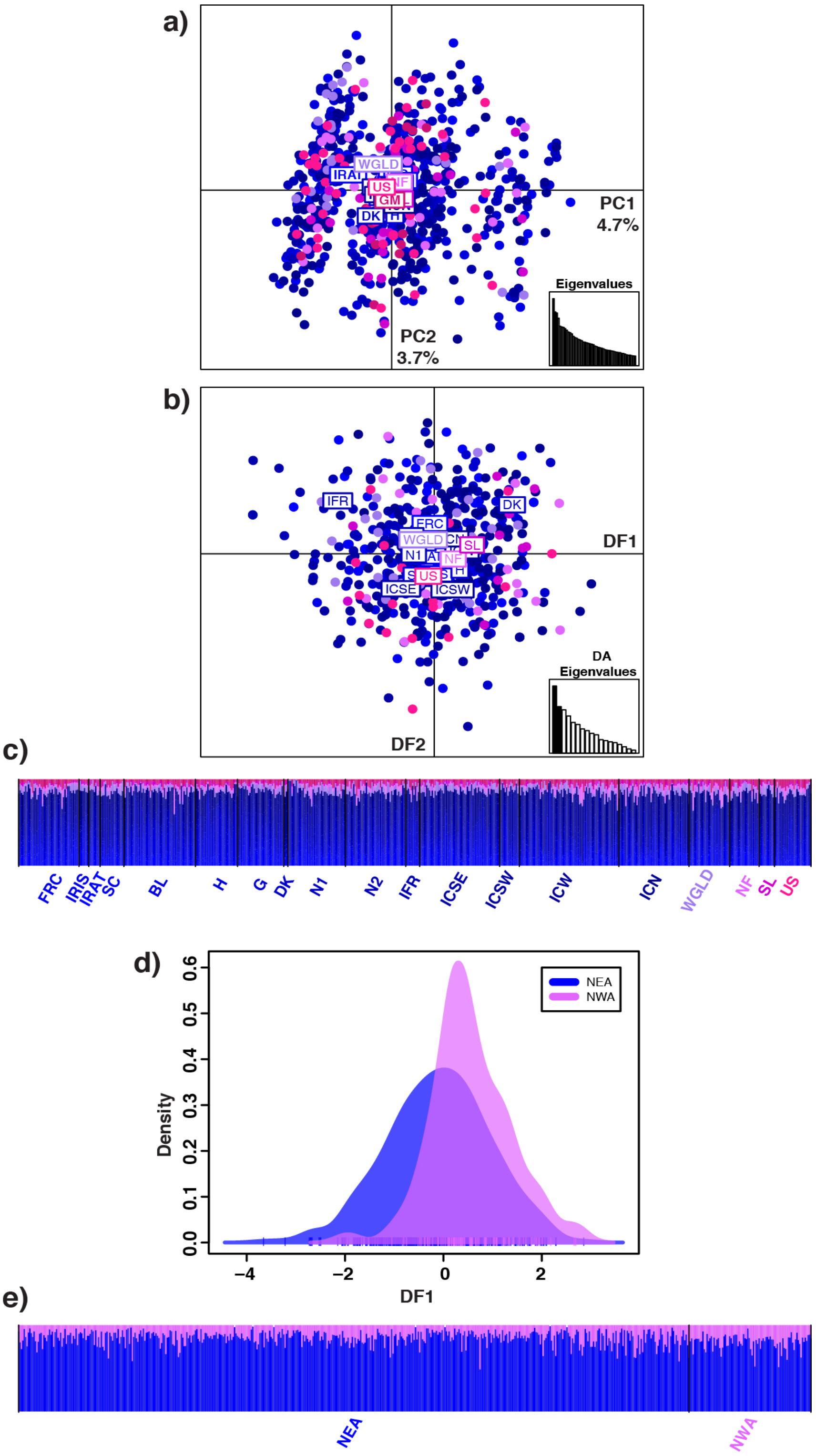

**Figure S8.** DAPCs focusing on the *P. p. phocena* subspecies in the North Atlantic. (a) PCA scatter plot showing the two first PCs. (b) DAPC scatter plot optimizing the difference among geographical subgroups showing the two first DFs. (c) Barplot showing the cluster membership probability of the DAPC presented in (b). (d) DAPC density plot optimizing the difference West and East individuals showing the two first DFs. (e) Barplot showing the cluster membership probability of the DAPC presented in (d). Acronyms are presented in Table S2. NEA=North East Atlantic. NWA=North West Atlantic.

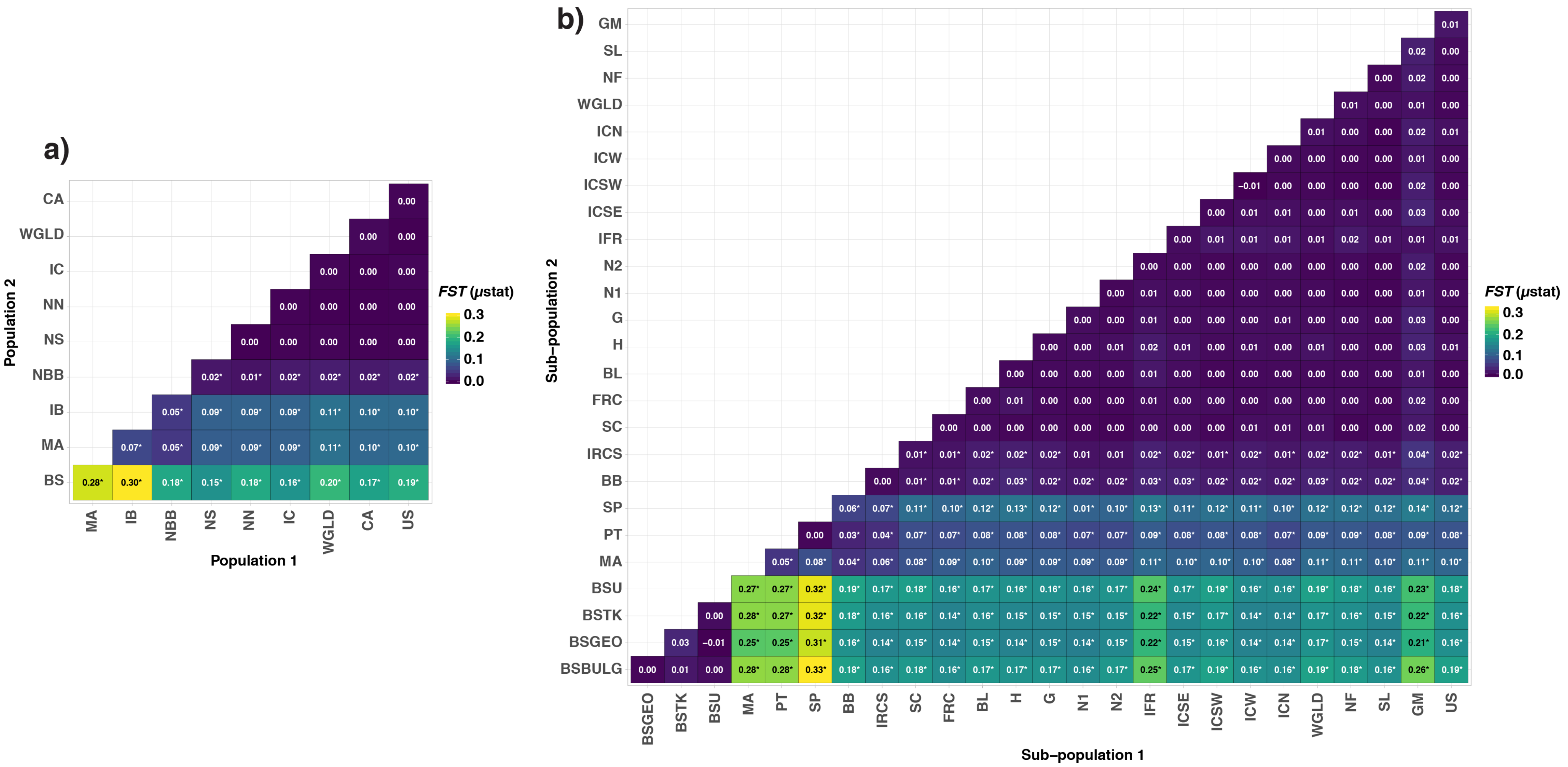

**Figure S9.** Pairwise $F_{ST}$ values between geographical regions (a) or subgroups (b) for microsatellites. Acronyms are presented in Table S1 (a) and Table S2 (b). *$p$-value < 0.05

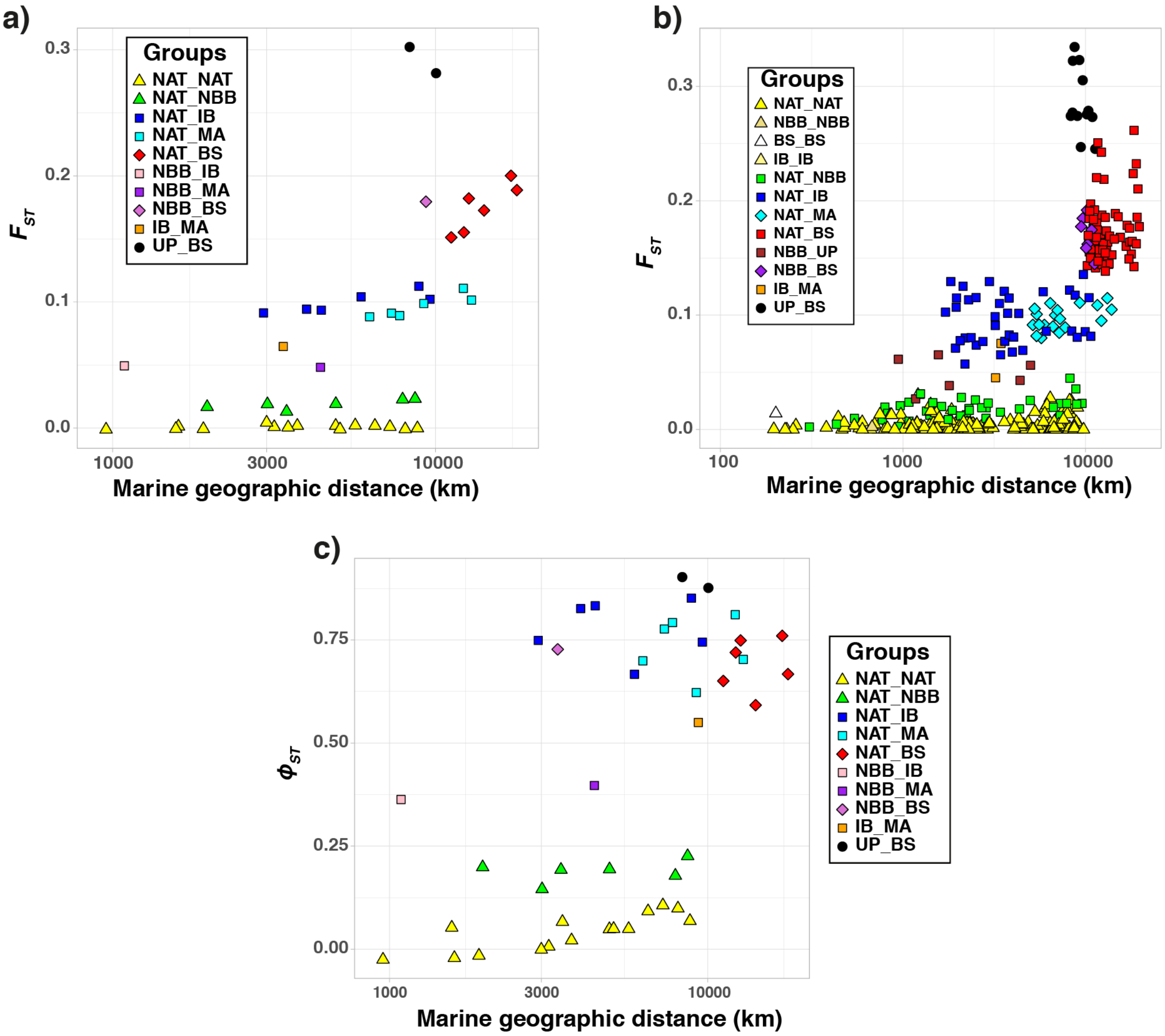

**Figure S10.** Relationships between geography and genetic distances. (a) Scatter plot of the genetic and marine geographic distance for all pairs of geographical regions using microsatellites. (b) Scatter plot of the genetic and marine geographic distance for all pairs of subgroups using microsatellites. (c) Scatter plot of the genetic and marine geographic distance for all pairs of geographical regions using mtDNA. UP=Upwelling. The other acronyms are presented in Table S1.

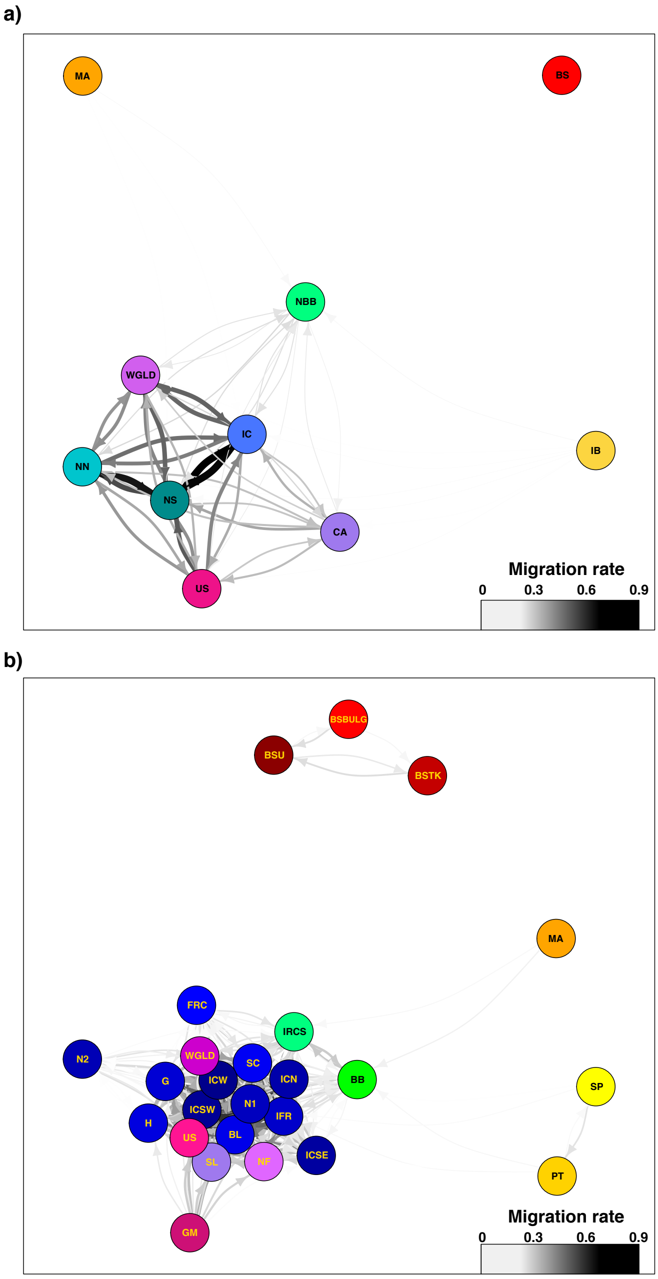

**Figure S11.** Genetic connectivity networks. Directional relative migrate rate (*m*) networks just above the percolation threshold among geographical regions (a) and subgroups (b). Acronyms are presented in Table S1 (a) or Table S2 (b).

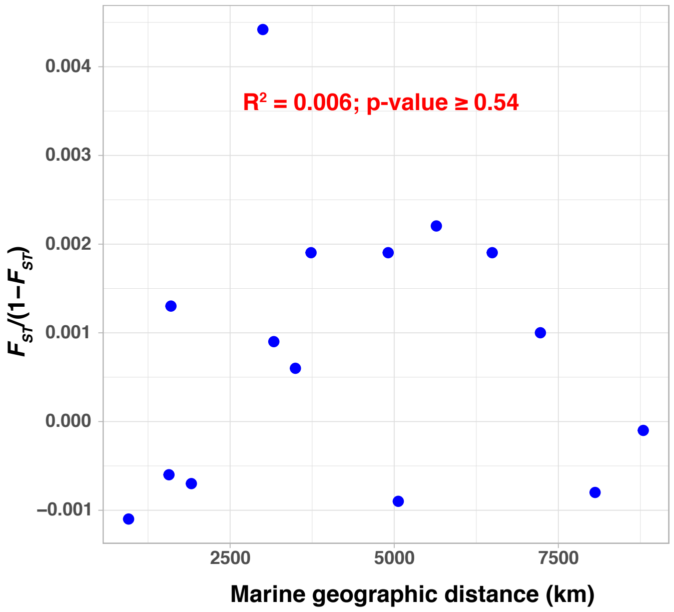

**Figure S12.** Pattern of isolation by distance in the North Atlantic. Scatter plot of the genetic and marine geographic distance for all pairs of geographical regions using microsatellites. The *p*-value was assessed using a Mantel test. The red line represents the regression line and $R^2$ represents the coefficient of determination.

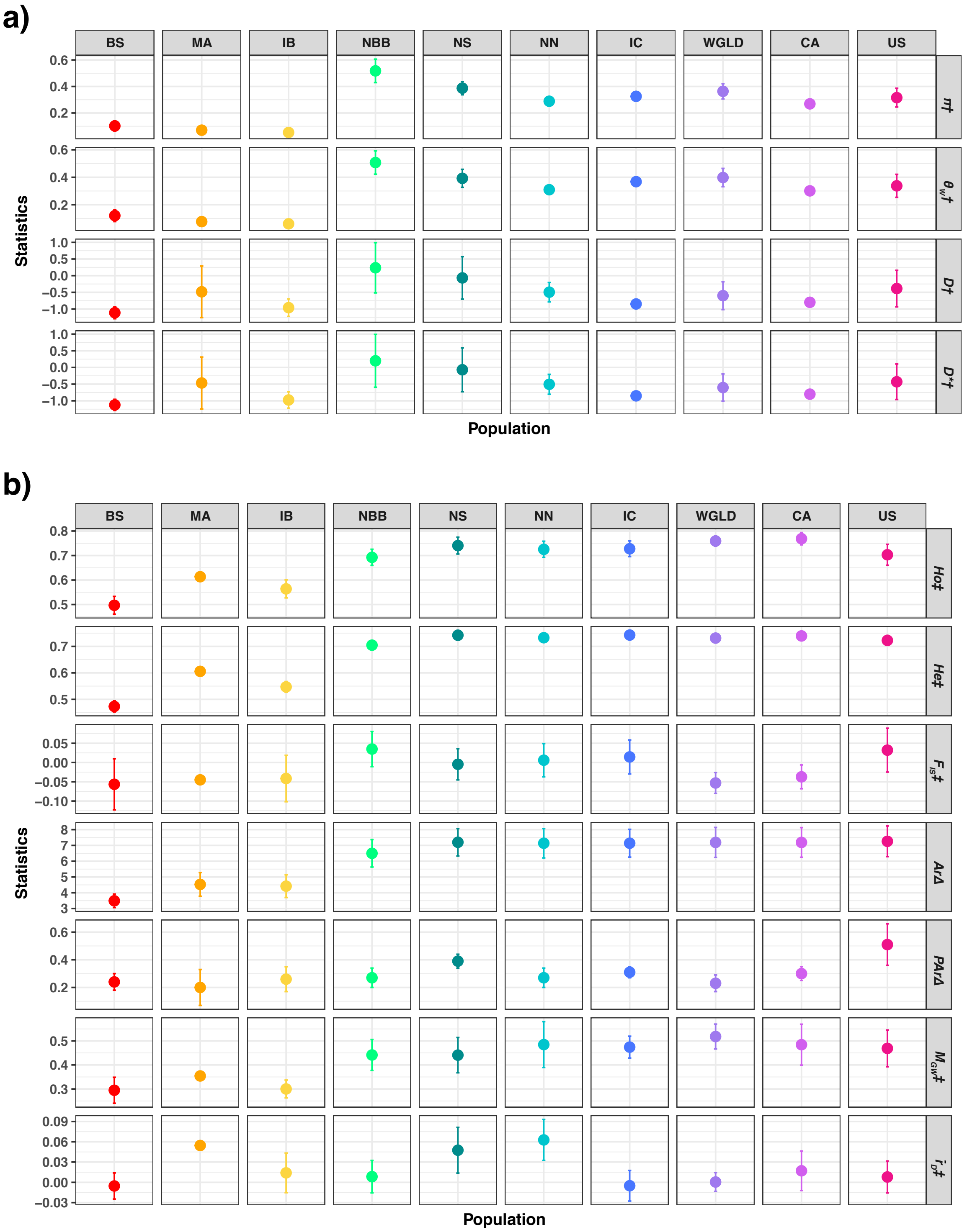

**Figure S13.** Genetic diversity statistics estimated among geographic regions for (a) mtDNA and (b) microsatellite loci. A rarefaction approach is used to account for difference in sample size among regions. (see Table 1 for the acronyms). $\pi$, nucleotide diversity; $\theta_W$, Watterson's theta; *D*, Tajima's *D*; *D**, Fu and Li's D*; *Ho* and *He*, observed and expected heterozygosity; $F_{IS}$, fixation index; *Ar*, Allelic richness; *PAr*, private allelic richness; $M_{GW}$, M-ratio; $r_D$, multilocus linkage disequilibrium statistic. Standardization assuming 5 (†), 14 (‡); and 18 gene copies (Δ). Acronyms are presented in Table S1.

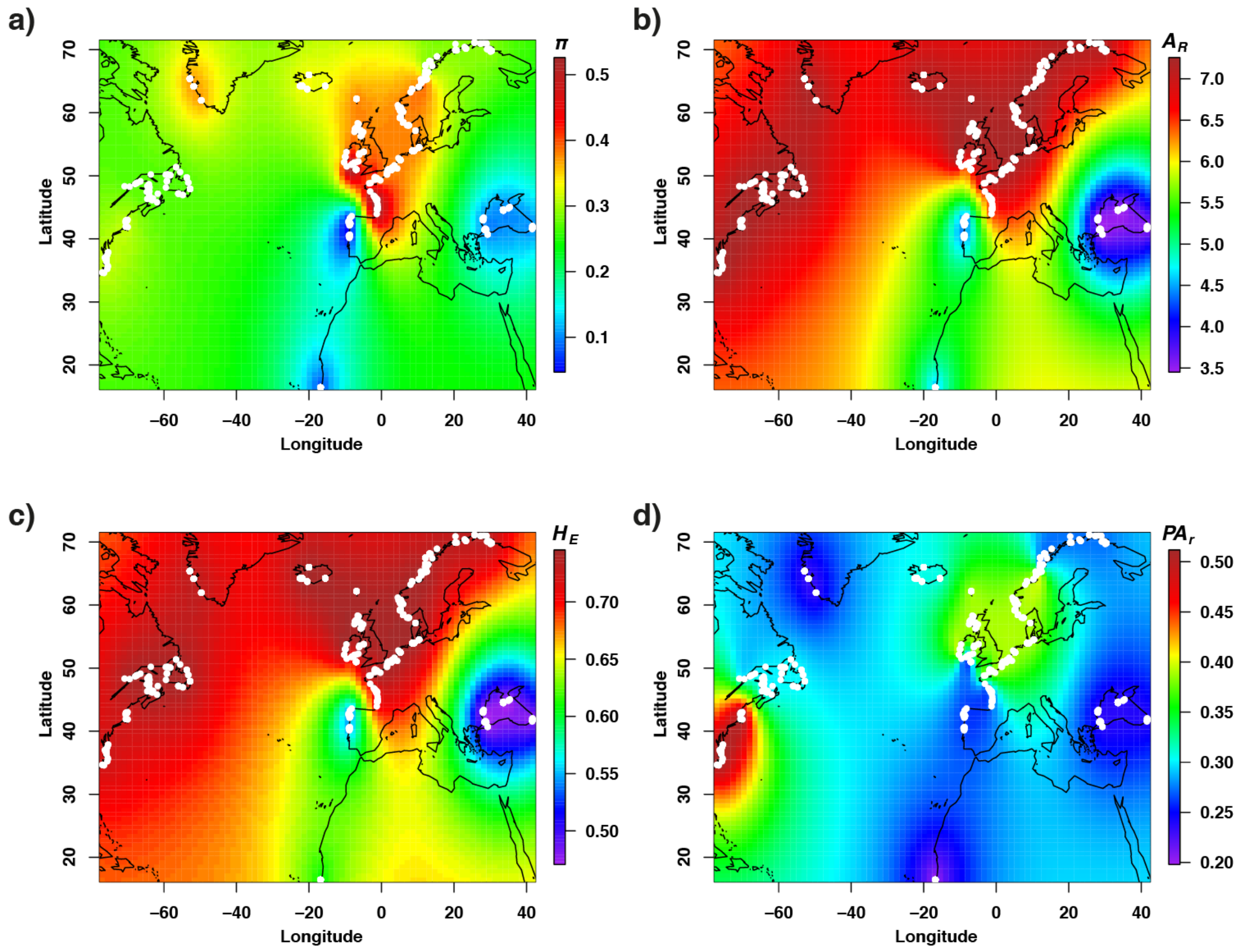

**Figure S14.** Spatial patterns in genetic diversity at the mitochondrial (a) and nuclear (b-d) level among the 10 geographical regions. (a) Average nucleotide diversity assuming a standardized sample size of 5 individuals per region. (b) Average allelic richness assuming a standardized sample size of 18 gene copies. (c) Average expected heterozygosity assuming a standardized sample size of 14 individuals. (d) Average private allelic richness assuming a standardized sample size of 18 gene copies. White dots indicate sampling points used to carry out the interpolation.

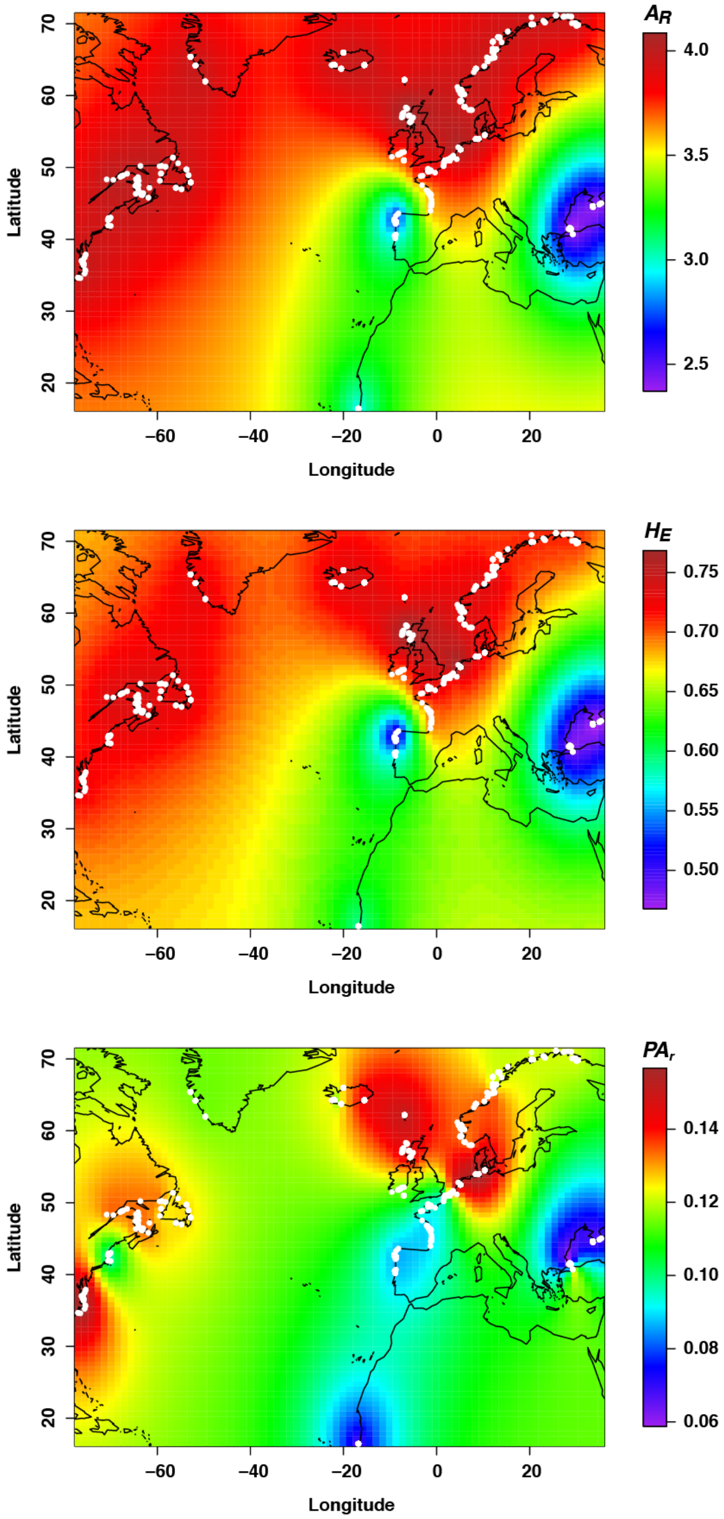

**Figure S15.** Spatial patterns in genetic diversity at the nuclear level among the 30 geographical subgroups. (a) Average allelic richness assuming a standardized sample size of 14 gene copies. (b) Average expected heterozygosity assuming a standardized sample size of 10 individuals. (c) Average private allelic richness assuming a standardized sample size of 14 gene copies. The white dots represent the point used to carry out the interpolation.

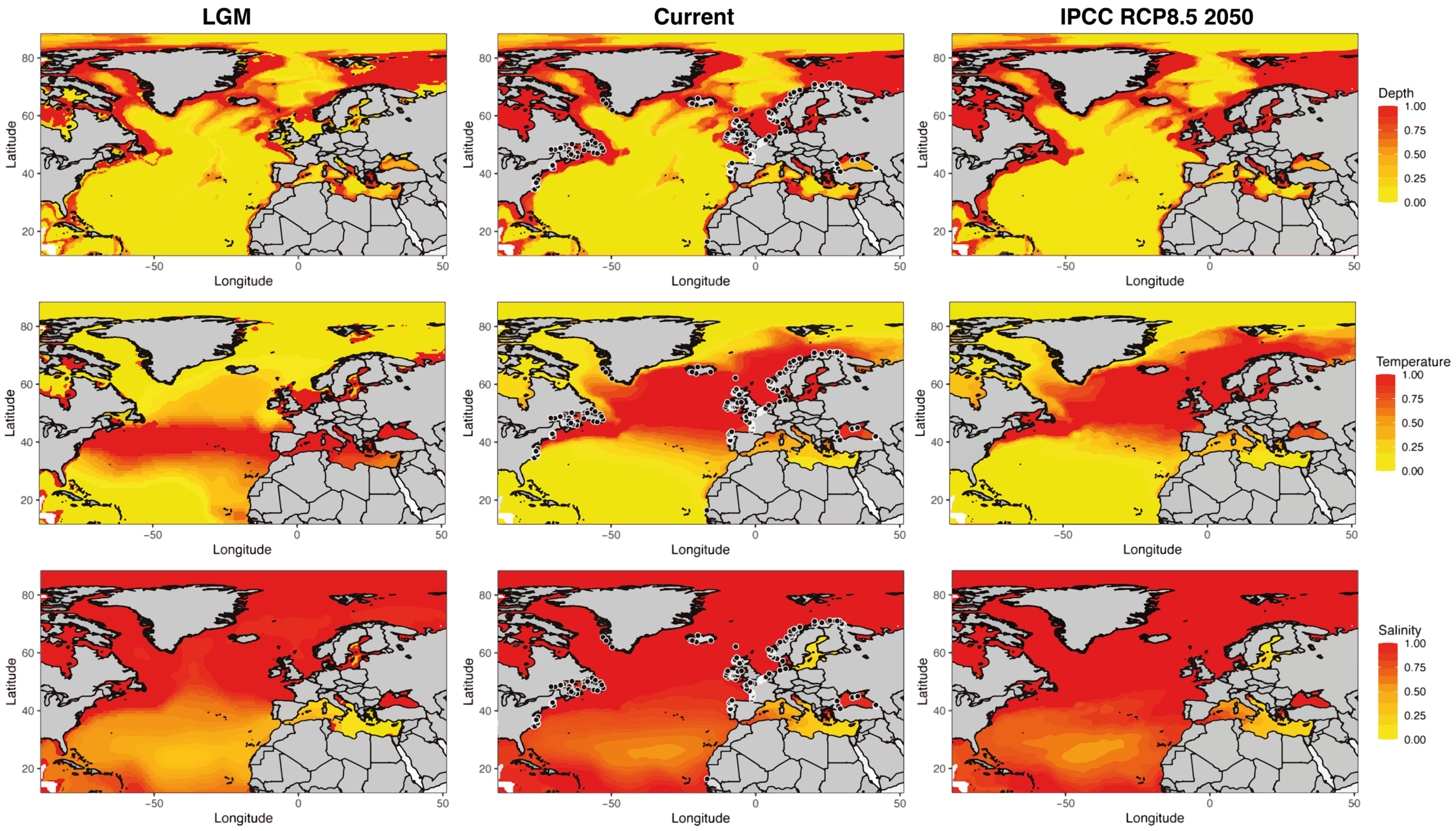

**Figure S16**. Suitability envelopes for each individual environmental variable used in AquaMaps to predict the global niche habitat suitability of harbor porpoises in the North Atlantic and adjacent seas for three time periods: last glacial maximum (LGM), current time and year 2050 under the RPC8.5 IPCC scenario. Black dots on the current maps show individual locations of the porpoise samples used in this study.

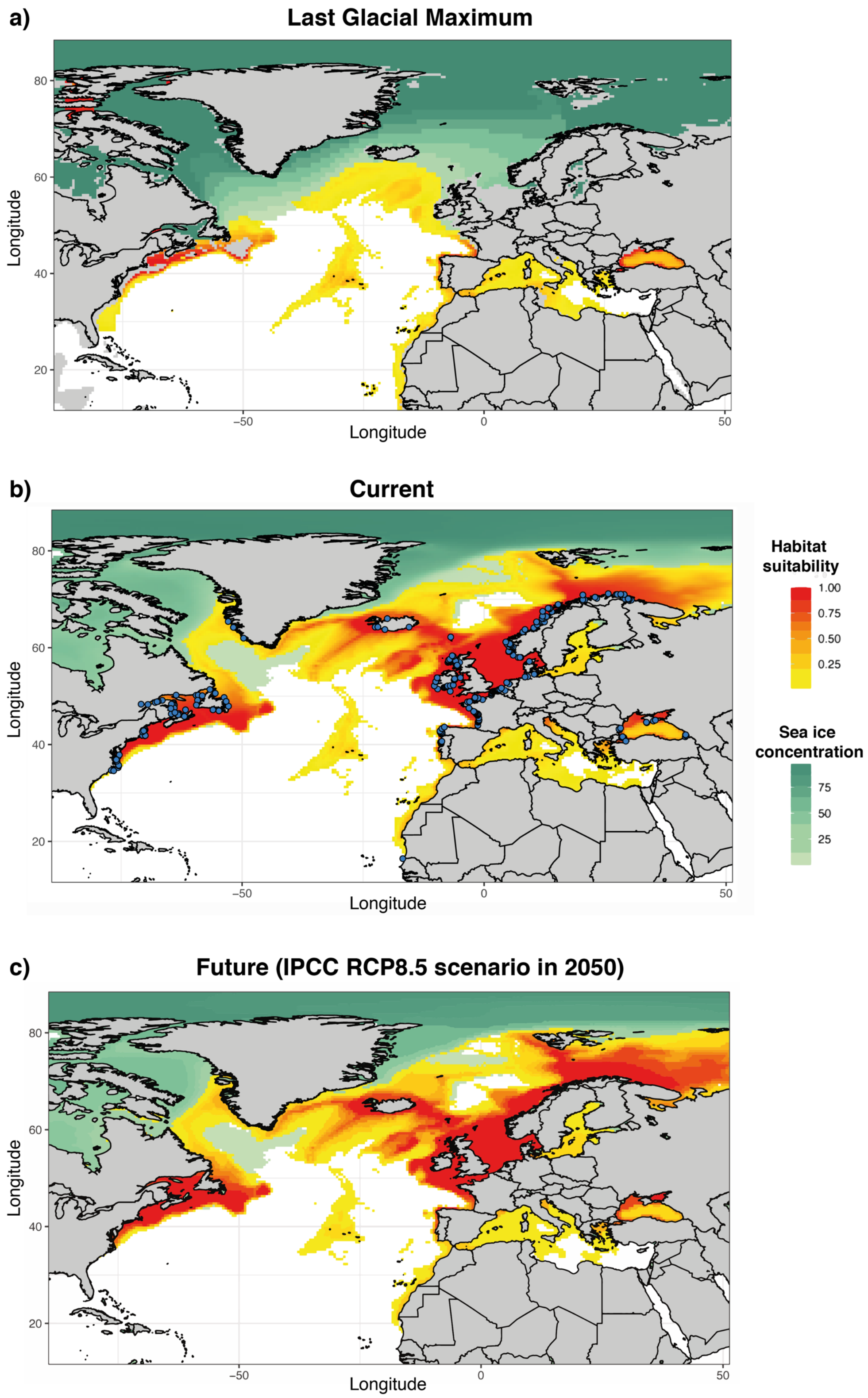

**Figure S17**. Maps showing the predicted habitat suitability for harbor porpoises throughout the North Atlantic and adjacent seas during three time periods generated using AquaMaps environmental niche modelling and input parameter settings described in Table S9, including salinity as additional predictor. Yellow to red colors represent least to most suitable habitat, respectively, based on the AquaMaps habitat model. Light to dark green colors represent the proportion of sea ice concentrations (%). Blue dots on the current map show individual locations of the porpoise samples used in this study.

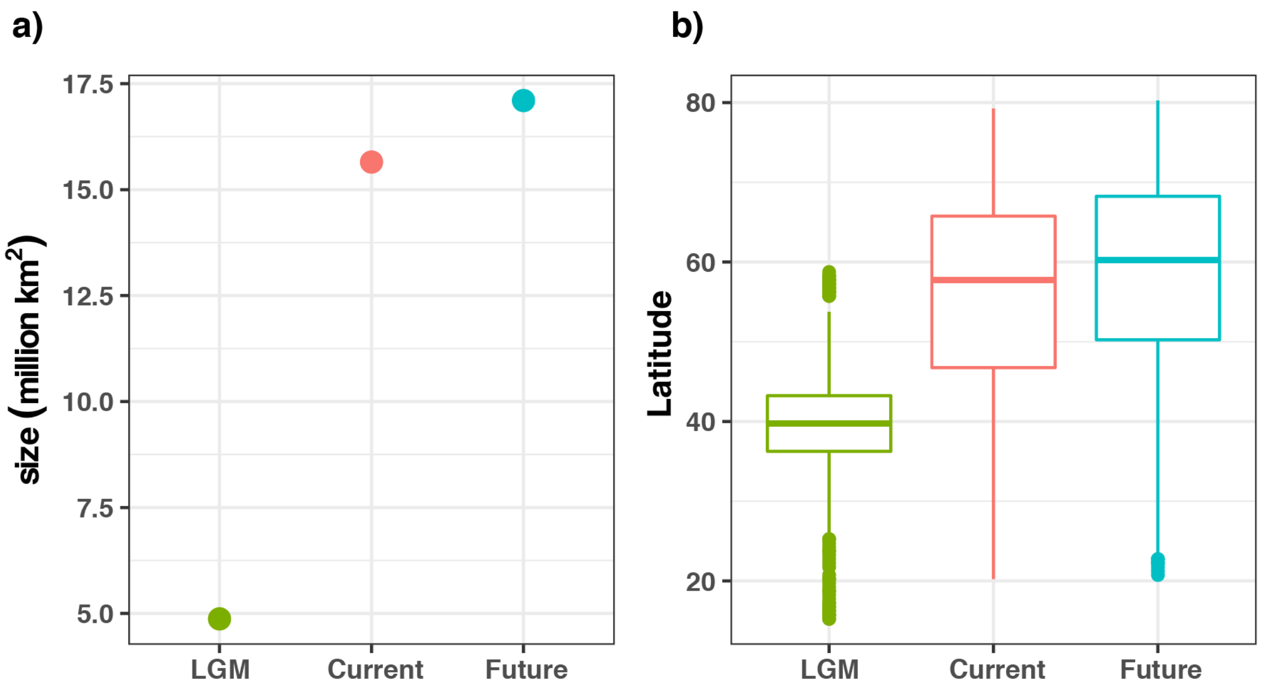

**Figure S18**. Habitat size (km$^2$) and average latitude of suitable habitats predicted by the AquaMaps relative environmental suitability ≥ 0.3 for harbor porpoises in the North Atlantic and adjacent seas.
